# Functional architecture for speed tuning in primary visual cortex of carnivores

**DOI:** 10.1101/2025.11.04.686504

**Authors:** Víctor Manuel Suárez Casanova, Noah K. Lasky-Nielson, Lihanxi Ye, Jonathan D. Touboul, Jérôme Ribot, Stephen D. Van Hooser

## Abstract

Perception of motion critically depends on detecting the speed and direction of moving stimuli. The primary visual cortex (V1) of some mammals, including primates and carnivores, exhibits functional organization for key receptive field properties such as orientation, direction, and spatial frequency; however, less is known about the organization of speed-tuned cells. While individual V1 neurons have been shown to exhibit speed selectivity, functional architecture for speed preference has been primarily reported in higher cortical areas such as primate area MT. Using multi-channel electrophysiology in anesthetized female ferrets, we investigated the joint tuning of V1 neurons for spatial frequency, temporal frequency, orientation/direction, and speed. We found significant clustering of cells tuned for speed and for speed preference within single electrode penetrations. We found that both simple and complex cells can exhibit speed tuning, and no strong variation across cortical layers. In reanalysis of intrinsic signal imaging data from cat V1, we observed repeating “hot spots” of high speed selectivity separated by “cold spots” with low tuning for speed. These findings indicate that a functional architecture for speed tuning is present within V1 itself and transmitted to downstream cortical regions.

**Significance Statement:** To perceive moving objects, the visual system must detect both their direction and speed. The primary visual cortex, the first cerebral visual area to receive visual information from the retina via the lateral geniculate nucleus, plays a key role in this process. Here, we demonstrate that the primary visual cortex in carnivores contains speed-tuned neurons. Moreover, these neurons are organized into clustered “hot spots” that repeat across the cortical surface, suggesting a functional architecture for speed. While speed-tuned functional maps were previously thought to exist only in higher visual areas, such as area MT in primates, our findings reveal their presence at the level of primary visual cortex.

## Introduction

Sensory processing is fundamental for animals to navigate their environment and respond to external stimuli, such as locating prey or avoiding predators. To process the movement of objects in an animal’s visual environment, the visual system must interpret, decode, and encode various stimulus features such as direction and speed. In primates and carnivores, specific regions of the primary visual cortex (V1) have evolved to exhibit a highly organized functional architecture, where neurons are tuned to various parameters such as orientation, direction, and spatial frequency (SF) (Hubel and Wiesel, 1959, 1962, 1965, 1968, 1997, 1998; Hawken et al., 1988; Bonhoeffer and Grinvald, 1991; Snodderly and Gur, 1995; Ohki et al., 2005).

The organization of these properties within V1 has been studied extensively, but how these parameters interact to encode more complex visual attributes, such as speed, remains unclear. Early studies posited that speed information is primarily processed in higher-ordered visual areas such as the medial temporal (MT) area in primates and the posteromedial lateral suprasylvian (PMLS) area in felines (Born and Bradley, 2005; Priebe et al., 2006 p.200; Zeki, 2015). However, recent findings show that V1 may also encode speed information (Orban et al., 1986a; Baker, 1990; Moore, 2006; Perrone, 2006; Price et al., 2006; Priebe et al., 2006). Additionally, studies have shown that direction preference in V1 often varies with stimulus temporal frequency (Moore et al., 2005; Cosnier-Horeau et al., 2024), suggesting that V1’s representation of motion is more complex: in particular, this data indicates the presence of a co-representation of motion with other stimulus features.

To further understand motion processing in V1, we characterized the joint tuning of neurons in ferret (Mustela putorius furo), whose visual system is well described and shares similar cortical organization to other carnivores and to primates (Bonhoeffer and Grinvald, 1991; Chapman and Stryker, 1993; Weliky and Katz, 1994; Chapman et al., 1996; Hübener et al., 1997; Rao et al., 1997; Issa et al., 1999; Katz and Crowley, 2002; Yacoub et al., 2007, 2008; Smith et al., 2015; Gilardi and Kalebic, 2021). Using *in vivo* electrophysiology, we examined tuning responses of V1 cells across layers with a multichannel silicon electrode (NeuroNexus) during visual stimulation with drifting sinusoidal gratings with several combinations of orientations, spatial frequencies, and temporal frequencies. Our findings demonstrate that simple and complex cells in ferret V1 can exhibit speed tuning, with no substantial difference across cortical layers. Interestingly, the speed-tuned cells display clustering into “hot” (high speed selectivity) and “cold” (low speed selectivity) zones. A reanalysis of intrinsic signal imaging data collected from cats (Sauvage et al., 2017; Cosnier-Horeau et al., 2024) revealed a functional map with periodicity similar to direction preference, arguing that zones of high and low speed tuning are part of each hypercolumn. These findings indicate that functional architecture for speed tuning emerges at the earliest cortical stage before being further elaborated in higher visual areas.

## Materials and Methods

All experimental procedures were approved by the Brandeis University Animal Care and Use Committee and performed in compliance with the National Institutes of Health guidelines.

### Animal Sourcing and Housing

Adult female (jill) ferrets (*Mustela putorius furo*; n = 16, age > 1 year) were obtained from Marshall Bio-Resources and used in terminal electrophysiological experiments designed to record activity across layers in the primary visual cortex (V1). Animals were housed in a humidity- and temperature-controlled room and entrained to a 12-h light-dark cycle via timed lights in a custom stainless-steel cage (60 cm x 60cm x 35 cm) equipped with a hammock and small toys. Animals had *ad libitum* access to food and water. Female animals were used exclusively due to co-housing with males resulting in undue distress in sexually mature jills.

### Surgical Procedures

Ferrets were initially sedated using ketamine (20 mg/kg i.m.). Atropine (0.16-0.8 mg/kg i.m.) and dexamethasone (0.5 mg/kg i.m.) were administered to reduce salivary and bronchial secretions and reduce inflammation, respectively. Animals were anesthetized using a mixture of isoflurane, oxygen, and nitrous oxide delivered through a mask while a tracheostomy was performed. After completion of the tracheostomy, animals were ventilated with a 1.5 % - 3 % isoflurane in a 2:1 ratio of nitrous oxide to oxygen, respectively. A cannula was inserted into the intraperitoneal cavity for delivery of 5 % dextrose in lactated Ringer’s solution (3 ml/kg/h i.p.). Body temperature was maintained at 37 °C using a thermostatically controlled heating pad. End-tidal CO2 levels and respiration rate were monitored and adjusted to maintain an appropriate physiological range (3.5% - 4%). The animal was secured in a custom stereotaxic frame that did not obstruct vision and silicon oil was applied to the eyes to prevent corneal damage. All wound margins were infused with bupivacaine (0.1 ml per wound site, 2.5 mg/mL). A midline incision was made on the scalp and the skull was exposed. A craniotomy (4 x 4 mm) was made on the right hemisphere and the durotomy was performed using a 31-gauge needle and dura pick. Prior to beginning electrophysiological recordings, the animals were paralyzed using the neuromuscular blocker gallamine triethiodide (10 - 30 mg/kg/h) delivered through the intraperitoneal cannula to suppress spontaneous eye movements, and the nitrous oxide-oxygen mixture was adjusted to 1:1. The animals’ ECGs were continuously monitored to ensure adequate anesthesia, and the percentage isoflurane was increased if the ECG indicated any distress.

### Electrophysiology

Electrophysiological recordings employed 32-channel single-shank laminar probes (NeuroNexus, A1x32-Poly2-5mm-50s-177-A32, 50 μm thickness) to record from several layers of the ferret primary visual cortex simultaneously (Ritter et al., 2017). The probe was positioned approximately perpendicular to the surface of the cortex and lowered until all pads were inserted into the brain (900 - 1100 μm), and 2% agarose was applied to prevent brain pulsation. Mineral oil was applied to the agarose at regular intervals to prevent the agarose from drying. An adaptor (NeuroNexus, A32-OM32) connected the electrode to a head stage (Intan Technologies, RHD2132). Signals were amplified using an RHD2000 amplifying/digitizing chip and USB interface board (Demo Board, Intan Technologies) and stimulus information was acquired using a Micro1401 acquisition board and Spike2 software (Cambridge Electronic Design).

Individual spike waveforms were extracted using JRCLUST (Jun et al., 2017) using 5 SDs as a threshold (on any channel) and data from all channels were used to define the spikes. The spikes were clustered by JRCLUST’s automated routine and then manually refined and classified by the experimenter as “single units” when waveforms appeared to reflect a single underlying process or “multi-units” when waveforms appeared to reflect multiple underlying processes.

### Visual Stimulation

Visual stimuli were created in MATLAB (MathWorks) with the Psychophysics Toolbox (Brainard, 1997; Pelli, 1997) and displayed on a 21-inch flat face CRT monitor (Sony, GDM-520) with a resolution of 800 x 600 pixels and a refresh rate of 100 Hz. The monitor was placed at a distance (20 - 40 cm) in front of the ferret such that it subtended 37.3 x 63.8 degrees of visual space. Receptive fields were manually mapped by displaying circular patches of drifting sinusoidal gratings at different positions and moving the monitor to accommodate different eccentricities while listening to the responses on a loudspeaker. All stimuli described below were developed in-house and are available at https://github.com/VH-lab/vhlab-NewStim-matlab.

### Co-Varied Joint Tuning Stimuli

Drifting grating visual stimuli were full-field, high contrast sinusoidal gratings (1 sec stimulation, 1 sec ISI) presented pseudo-randomly. Each individual grating stimulus had a fixed direction, spatial frequency, and temporal frequency, and these parameters were co-varied within a set so that all combinations of orientation or direction, spatial frequency, and temporal frequency were shown in the same recording epoch. A total of 8 directions of motion (in steps of 45°), 8 different spatial frequencies (0.04 - 1.25 cycles per degree) and 7 temporal frequencies (0.5 - 32 Hz) were presented at 100 % contrast. Therefore, a total of 448 stimuli were presented, each repeated 5 times. In our first 12 experiments, gratings at a particular orientation drifted back and forth allowing us to assess responses to stimulus orientation. In our last 8 experiments, stimuli drifted in a single direction, allowing us to assess direction selectivity.

### Blinking Stimuli

A full-field blinking stimulus was employed to measure current source density (Stitt et al., 2013). The stimulus was presented in full-field view of the animal with a 100 ms ON and 200 ms OFF interval for 50 repetitions.

### Intrinsic signal imaging

We re-analyzed data taken from 3 isoflurane-anesthetized cats (*Felis catus*) from previous work (Ribot et al., 2016; Sauvage et al., 2017; Cosnier-Horeau et al., 2024). In brief, all combinations of spatial frequency (0.15, 0.29, 0.5, 0.9, and 1.65 cpd cycles/degree), temporal frequency (0.38, 0.88, 1.24, 1.53, 1.65, 2, 2.78, 3.26, 3.85, 5.37 Hz), and direction tuning were examined. This was achieved by showing a drifting grating at a given spatial and temporal frequency, and slowly changing the angle of drift (stimulus direction) counter-clockwise at a rate of 1/60 Hz. Orientation tuning was then inferred from the phase at the second harmonic of the Fourier transform of the hemodynamic signals, as described in (Kalatsky and Stryker, 2003). Direction tuning estimated by inverting the filtering associated with hemodynamic signals as described in (Sauvage et al., 2017), and data were discretized into 200 bins around the full circle 360°.

### Data and Statistical Analysis

Data were analyzed in MATLAB using the Neuroscience Data Interface (García Murillo et al., 2022). A total number of 458 V1 neurons were identified via spike sorting. For each neuron, the mean response (F0) and modulation at the stimulus frequency to drifting grating stimulation (F1) were examined. If a cell’s F1 response was greater than the mean response (F0), F1 was used to calculate index values. If F0 was greater, it was used for calculations (Movshon et al., 1978a, 1978b, 1978c; Heimel et al., 2005). Sites were only included in the analysis if they exhibited significant variation across all stimuli by an ANOVA test, p < 0.05. All recordings in each experiment had at least 7 pass these exclusion criteria.

### Correlations

For electrophysiology, correlations were computed using Pearson’s correlation (Matlab function: corcoeff) and significance evaluated with a custom Matlab function (vlt.stats.corrcoefResample) (**github.com:VH-Lab/vhlab-toolbox-matlab**) that computes the correlation coefficient for random shufflings of the X and Y pairs to be considered and finds where in the distribution of shuffled correlation coefficient values the actual data lies.

To determine the statistical significance of correlations between the tuning properties of pixels in intrinsic signal imaging, it was necessary to control for the fact that the samples were not independent and that each individual property being examined (speed index, orientation selectivity index, etc) had spatial dependence. To evaluate the significance of the correlation coefficient calculated with the actual data, we calculated the correlation coefficient in 10,000 surrogate datasets where one of the maps was randomly shifted in x and y. The shift was a toroidal shift, so that the portion of the image that ran off the edges of the original image was wrapped around to the other sides. We used the 10,000 surrogate datasets as the expected distribution of values between two maps with the spatial autocorrelations that were inherent to each map, and found the percentile value of where the correlation coefficient of the actual data fell in this distribution. We made a custom function vlt.stats.spatialCorrelationSignificance (**github.com:VH-Lab/vhlab-toolbox-matlab**) for this purpose.

### Speed Tuning

We examined responses to the co-varied stimuli and determined the preferred orientation (early experiments) or direction (later experiments) by finding the orientation/direction of the stimulus that produced the maximum response (in either F0 or F1). Then, we fit responses to this orientation or direction for all combinations of temporal frequency/spatial frequency that were fit with a product of Gaussians function as in Priebe et al. (2006):

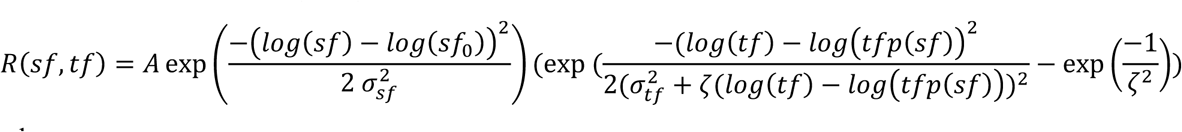

where

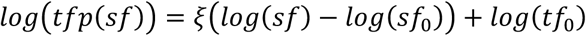

In this context, *A* represents the neuron’s peak response,*sf*_0_ denotes the preferred spatial frequency averaged over all temporal frequencies, and *tf*_0_ the preferred temporal frequency averaged over all spatial frequencies. The function tfp(sf) allows the temporal frequency to have a dependency (ξ, the speed index) on the spatial frequency. The variable ζ characterizes the skewness of the temporal frequency tuning curve.

For both electrophysiology and intrinsic signal imaging analyses, we excluded any fit that had an r^2^ value less than 0.2. In intrinsic signal imaging analyses, these pixels are shown in gray (same color as the background outside of the region-of-interest).

In many analyses, we examined whether there was significant evidence of speed tuning by using a nested fit analysis (Nested F test). We compared the squared error of the function above with a simpler function where the speed tuning parameter ξ was required to be 0 (our nested model). We computed the F statistic:

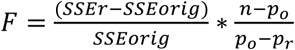

Where SSE_r_ and SSE_orig_ are the sum of squared errors for the nested and original models, respectively, p_o_ and p_r_ are the number of parameters for the original and nested models (7 and 6 here), and n is the number of datapoints used to generate the fit. The number of degrees of freedom in the F distribution are (*p_o_* − *p_r_*, *n* − *p_o_*). A p-value less than 0.05 indicated significant speed tuning.

### Orientation and Direction Tuning

We assessed orientation and direction tuning by analyzing responses to stimuli at the preferred spatial and temporal frequencies. To quantify selectivity, we calculated circular variance (CV) and direction circular variance (DCV) using fit-less vector measures as previously described (Ringach et al., 2002; Mazurek et al., 2014; Ritter et al., 2017; Stacy et al., 2023). CV was calculated from orientation tuning curves by measuring the mean spike rates, Rk, in response to a grating drifting at angle, *θ*_k_, which spanned from 0° to 360° in equally spaced intervals. C*V* approaches 1 for weakly selective cells and 0 for highly selective cells. The same method was applied for direction tuning, using directional circular variance (DCV), where angles represent directions in radians. These metrics provide a robust assessment of orientation and direction selectivity where higher values of 1 − *CV* and 1 − *DCV* indicate stronger selectivity:

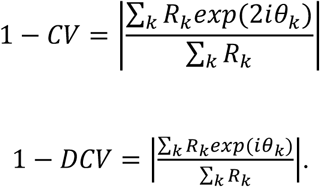

### Electrophysiology cluster analysis

To investigate the presence of speed-selective clusters in V1, we analyzed the speed index (SI) of cell pairs by comparing all pairs of cells from the same penetration or from different electrode penetrations. We categorized each pair based on whether both cells exhibited speed index values were both more or less than the median speed index (consistent with columnar organization) or if they were mixed (inconsistent with columnar organization). A similar analysis was conducted for orientation preference to confirm the known columnar organization in V1. In that analysis, we asked whether the pair’s orientation preferences were similar (closer than 45°) or dissimilar (not closer than 45°). To compute the significance for these pair analyses, we re-computed all values for 1000 random shufflings of the penetration numbers to which each cell was assigned and looked at how frequently we observed differences that were as great as the real data from which we estimated a p-value. In detail, given a pair within an electrode, we computed the fraction of pairs from the same electrode penetration that both exhibited speed tuning and subtracted the fraction of pairs from different electrode penetrations that both exhibited speed tuning, repeated the analysis for 1000 shuffles, and calculated where our actual data fell in the thus derived distribution to obtain a p-value.

### Laminar Analysis

To examine the relationship between speed tuning and cortical layer location, we first aligned recorded cells to standardized cortical depths using current source density (CSD) plots obtained during full-screen blinking stimuli presentations (Maier et al., 2010, 2011; van Kerkoerle et al., 2017). Layer 4 was identified based on the presence of a major current sink tightly locked to visual stimulus onset, allowing for the classification of cells into layers 2/3, 4, and 5/6. We then compared speed index values across these layers. Additionally, we assessed orientation tuning strength (1-CV) across the same layers to investigate any differences in tuning properties relative to cortical depth.

### Spatial analysis in intrinsic signal imaging data

To examine periodicity in speed tuning and direction selectivity, we used two measures, both performed on binary images where a nested F test provided evidence of significant speed tuning (significant speed index values) or where direction preference was determined to be within −30 to 30 degrees (direction control). First, we computed an autocorrelation map by computing the correlation coefficient between pixels at a given location and pixels located at shifts Δx and Δy away from the reference location.

In a second measure, we computed a 1-D power spectral density defined as the radial average of the usual 2-dimensional power spectral density (custom Matlab function (vlt.image.spatialPowerSpectralDensity). The image was padded on all sides with zeros to increase reporting resolution at low spatial frequencies and filtered with a Hanning window to reduce artifacts of the rectangular imaging window. In order to compute significance of power spectral density values that were measured, we created 1000 surrogate datasets where the locations of pixels that were inside the region of interest and had r2 values less than 0.2 were shuffled. We then computed the spectral power of these datasets including 95% confidence ranges and plotted them with the power spectral density values of the data. These shuffles controlled for the aperture of the imaging window, the region of interest, and the mean number of pixels that met the criteria.

### Histology

Prior to the start of recording experiments, the posterior surface of the multi-channel single shank electrode (NeuroNexus) was painted using fluorescent Di-I (DiCarlo et al., 1996; Liu, 2019). After the experiments, the electrode was withdrawn, and the ferrets removed from the custom stereotaxic frame. The animals were then administered pentobarbital sodium and phenytoin sodium (200-400 mg/kg, 25-50 mg/kg, respectively) intraperitoneally and prepared for perfusion. Animals were transcardially perfused, and the brain was fixed in 4% paraformaldehyde in 0.1 M PBS at 4 °C for 24 hours then moved to 30% sucrose in PBS for 48 hours. The recording hemisphere was sectioned sagittally into 100 μm sections using a sliding microtome (Leica SM2010R). Floating sections were washed in 0.1M PBS for 3×15 minutes and permeabilized in a blocking buffer (5% goat serum, 3% BSA, 0.3% Triton-X 100 in PBS) for 2 hours at room temperature. Sections were incubated with fluorophore-conjugated anti-NeuN antibodies (Alexa Fluor 488 Rabbit anti-NeuN, ab190195) at 1:300 dilution in the blocking buffer. Sections were then washed as previously described and mounted on electrostatic slides to air-dry. Slides were then cover-slipped using ProLong Diamond Antifade Mountant with DAPI (ProLong Diamond Antifade Mountant, ThermoFisher Scientific, P36962) and sealed using clear nail polish (Color 109 Invisible, Sally Hansen). Fluorescent images were taken using a fluorescent microscope (Keyence BZ-X710) and the electrode tracks were reconstructed using the Di-I traces (Van Hooser et al., 2003; Stacy et al., 2023; Zheng et al., 2023).

## Results

In this study, we set out to characterize the relationship between spatiotemporal processing and cortical architecture across the surface and layers of carnivore V1. In anesthetized ferrets, we inserted silicon electrodes with electrode sites spanning 750 µm in depth and recorded extracellular electrophysiological responses (**Figure 1A**). During recording sessions, we presented a battery of visual stimuli (**Figure 1B**) to assess the co-relationships between spatial frequency, temporal frequency, orientation, and direction preferences. Stimulus parameters were co-varied, shown in random order and repeated five times (~1.5-hour recording sessions). Our goal was to collect and characterize responses across the cortical depths to sample multiple V1 layers per recording (**Figure 1C**). To confirm recording locations, we coated the silicon electrode with fluorescent DI-I and reconstructed penetrations using post-hoc histological staining (**Figure 1C**).

**Figure 1.**
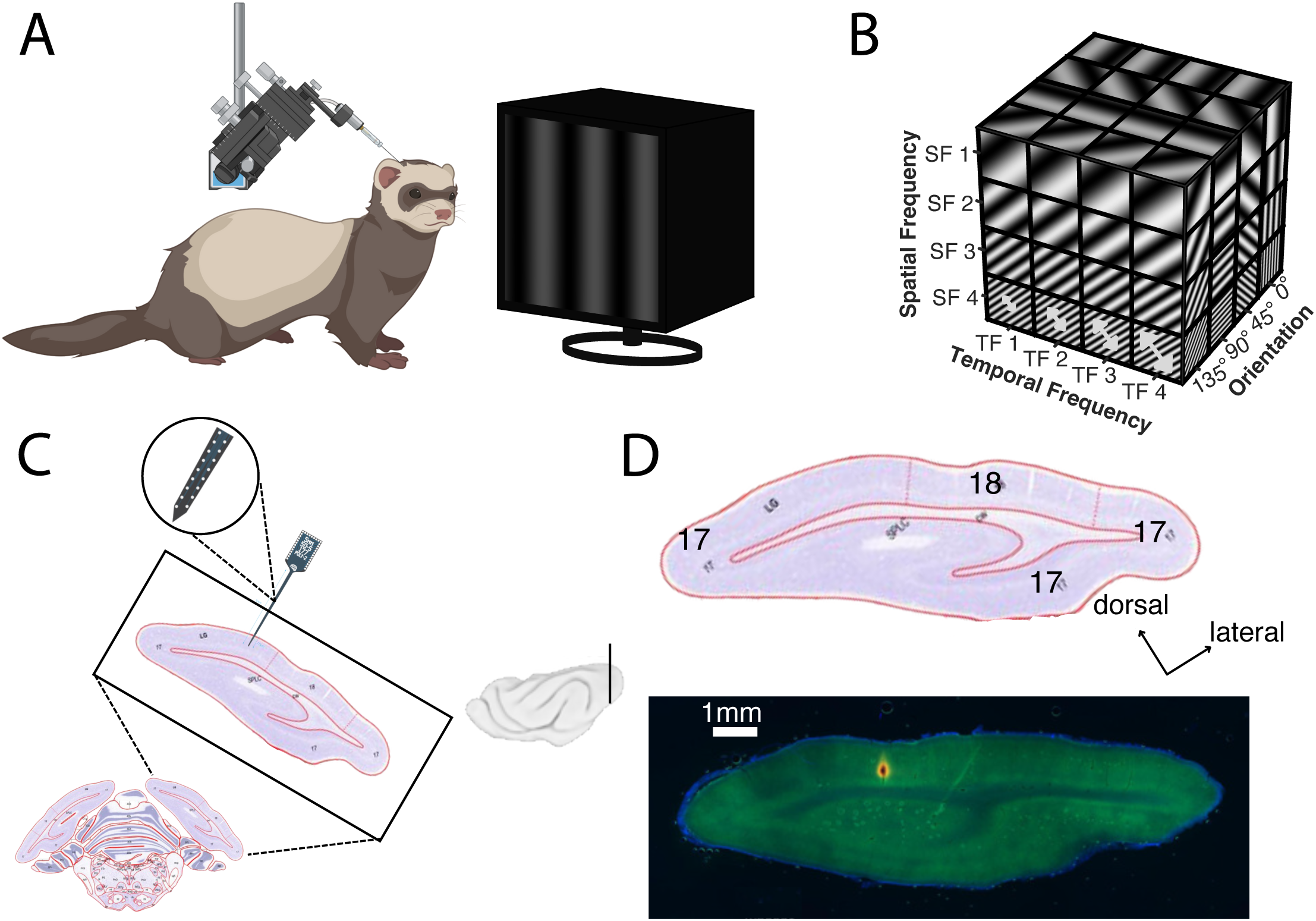
Experimental methodology. **(A)** We recorded evoked neuronal activity in anesthetized ferrets while presenting drifting grating stimuli. (B) Stimuli were drifting sinusoidal gratings with three parameters co-varied across all combinations: temporal frequency, spatial frequency, and direction. Arrows indicate increases in temporal frequency (back and forth motion in some experiments, single direction movement in other experiments); larger arrows indicate that more bars per second pass a given point (higher temporal frequency). These stimuli were presented in pseudorandom order. **(C)** NeuroNexus Electrode penetrations targeted V1; view from ferret atlas (Radtke-Schuller, 2018) shown. Plane of section is shown in brain image (inset) **(D)** Histological section shows penetration in V1. Electrode coated in Di-I (red); cortical neurons stained with NeuN (green); nuclei labeled with DAPI (blue). Images captured at 2X magnification.

### Speed tuning in the ferret visual cortex

The speed of a drifting grating is defined as its temporal frequency divided by its spatial frequency. Because of this relationship, a neuron cannot be independently tuned for to a specific spatial frequency and temporal frequency and also be tuned for speed. If a neuron responds best to the same speed across all spatial frequencies, then its preferred temporal frequency must increase proportionally with spatial frequency. In contrast, if the preferred temporal frequency remains constant as spatial frequency changes, the preferred speed must vary, indicating separable tuning for spatial and temporal frequency.

Following Priebe (et al. 2006), we visualized these tuning relationships using two approaches. First, we plotted speed tuning curves for each spatial frequency at the neuron’s preferred orientation or direction. For speed-tuned neurons, the curves align and peak at the same speed **(Figure 2A, left)**, whereas separably tuned neurons show shifting peaks **(Figure 2B, left)**. Second, we used heat maps of response amplitude across spatial and temporal frequencies. Speed-tuned neurons exhibit diagonal “slants” in these maps, reflecting temporal frequency changes in relation to spatial frequency (**Figure 2A, right**). In contrast, separable neurons exhibit rectangular or oval patterns without such slants (**Figure 2B, right**).

**Figure 2.**
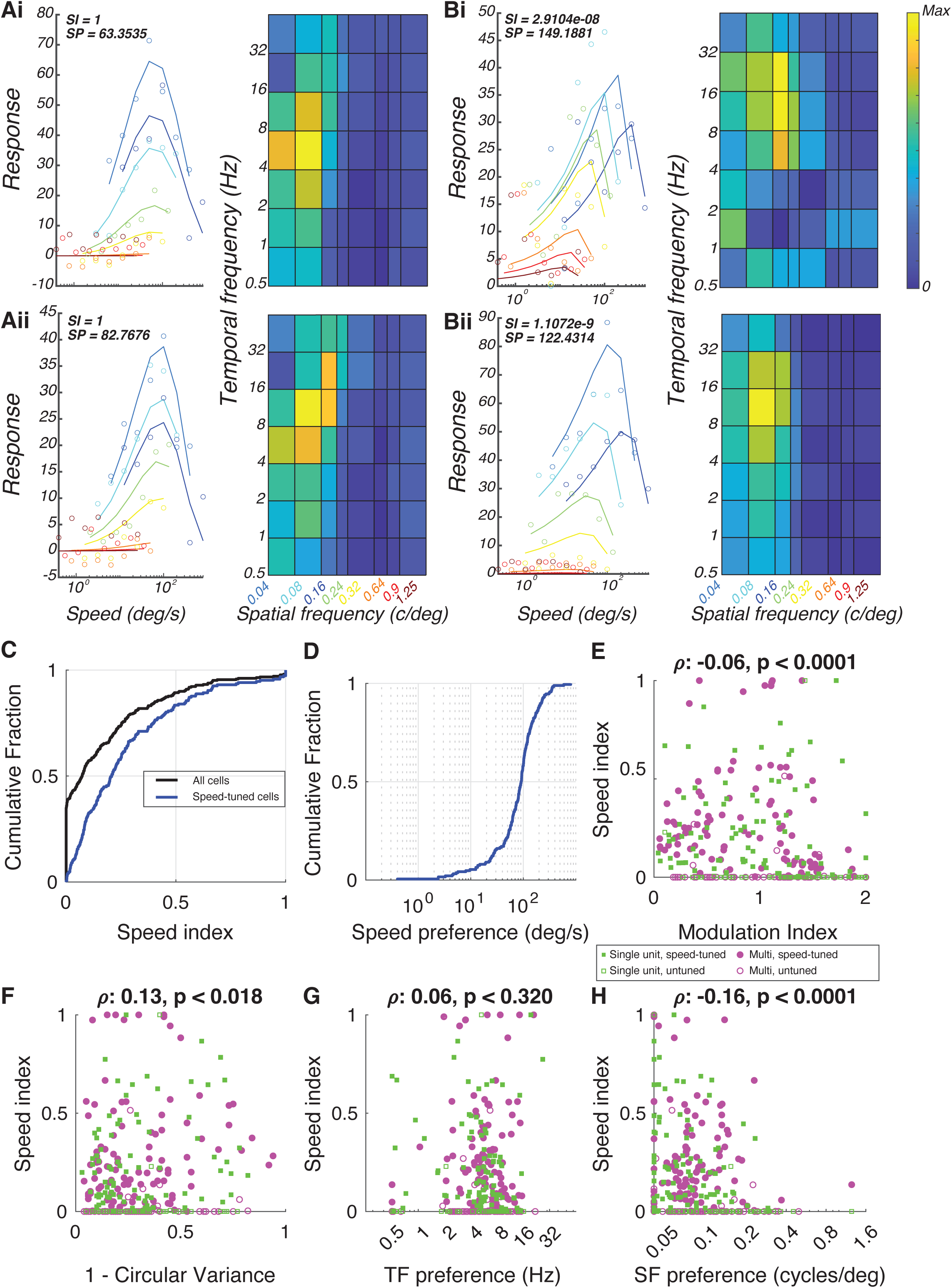
Speed tuning in individual neurons in ferret primary visual cortex. **(A)** Examples of speed-tuned cells in the primary visual cortex that exhibit high speed tuning (SI = 1), with varying speed preferences (SP = 63.35 deg/sec, top; SP = 82.77 deg/sec, bottom). **(B)** Speed-invariant cells were also identified across V1, with very low speed index values (SI ≈ 0). **(C)** A range of speed index values were observed, with about 40% of all cells (black) exhibiting an SI of 0. We calculated the distribution for cells that showed significant speed tuning by a nested F test (blue). **(D)** Significantly speed-tuned cells (blue) displayed a broad range of preferred speeds, from 4 to 1000 deg/sec, with most cells preferring speeds between 50 and 125 deg/sec. **(E)** Both simple and complex cells exhibited speed tuning, though complex cells were more likely to show higher SI values, indicating a negative correlation between SI and modulation index (single units: green squares; multi-units: magenta circles) (correlation coefficient, ρ = −0.06, p < 0.0001). **(F-H)** Correlation of SI with other tuning properties: SI and 1-CirVar **(F)** (correlation coefficient, ρ = 0.13, p = 0.018), spatial frequency **(G)** (correlation coefficient, ρ = 0.06, p = 0.320), and temporal frequency **(H)** (correlation coefficient, ρ = −0.16, p < 0.0001).

Across the population, speed index values ranged from 0 (no speed tuning) to >0.5 (strong speed tuning) (**Figure 2C**). The median cell exhibited relatively low speed tuning (0.073) but a minority of neurons (about 10%) exhibited speed index values greater than 0.5. To assess which cells were significantly speed-tuned, we applied a nested F test to compare models with and without the speed tuning parameter. This analysis revealed that 60.5% of cells were better fit by including speed tuning, and we classified these cells as significantly speed-tuned (Figure 2C). Among this subset, median speed tuning index values was 0.207. We then examined the preferred speed of cells with significant tuning (**Figure 2D**). Their preferred speed ranged from about 4 degrees per second to 1000 degrees per second, with most cells (25 - 75 percentile) exhibiting preferred speeds between 53 - 125 degrees per second.

We next investigated whether simple and complex cells could exhibit speed tuning, or if speed tuning was primarily confined to one of these cell types. To classify neurons as simple or complex, we used a modulation index calculation comparing the ratio between the stimulus-modulation response (F1) to the magnitude of the mean stimulus-evoked response (F0). Cells with F1/F0 ≥ 1 are defined as simple and those with F1/F0 <1 are defined as complex (Skottun et al., 1991; van Kleef et al., 2010; Wypych et al., 2012). We plotted the speed index value vs. the modulation index (**Figure 2E**). We observed a broad distribution of speed index values for both types. However, there was a significant negative correlation between speed index and modulation index (correlation coefficient, ρ = −0.06, t-test p = 0.0001), indicating that complex cells were more likely than simple cells to exhibit high-speed index values.

Finally, we examined whether speed tuning was correlated with other tuning properties. A weak but significant negative correlation (correlation coefficient, ρ = −0.13, shuffle p = 0.018) was found between speed index and orientation selectivity (1 – CV). There was a relatively weak relationship between speed index and temporal frequency preference (**Figure 2G**: correlation coefficient, ρ = 0.06, shuffle p<0.320) but there was a significant negative relationship between speed index and spatial frequency tuning preference (correlation coefficient, ρ = −0.16, p<1e-4). That is, cells with high speed index values were less likely to exhibit high spatial frequency values.

### Columnar clustering of speed-tuned cells and speed preference in cortical space

We next examined whether speed tuning in the ferret visual cortex follows a columnar organization, similar to that observed for other visual response properties such as orientation, spatial frequency, and temporal frequency. For each of the 20 electrode penetrations, we visualized individual neuron responses using “penetration plots” in which each cell’s speed index (SI) was plotted against its relative depth on the silicon probe, with 0 corresponding to the deepest recording site and 1 to the most superficial. **Figure 3** shows 12 of these electrode penetrations, ordered from left to right by increasing speed tuning (**Figure 3A**). In those on the left, most cells exhibited little or no speed tuning, whereas in those on the right, any neurons showed strong speed selectivity. These results suggested the presence of a columnar organization for speed tuning, with some columns containing many cells tuned for speed while others containing few such cells. To explore this further, we plotted preferred speed values for the subset of cells that passed the nested F test (**Figure 3B**). Within individual penetrations, preferred speed appeared strikingly similar across cells. Penetration plots for other tuning properties (spatial frequency preference, temporal frequency preference, orientation preference) are shown in **Supplementary Figure 1**.

**Figure 3.**
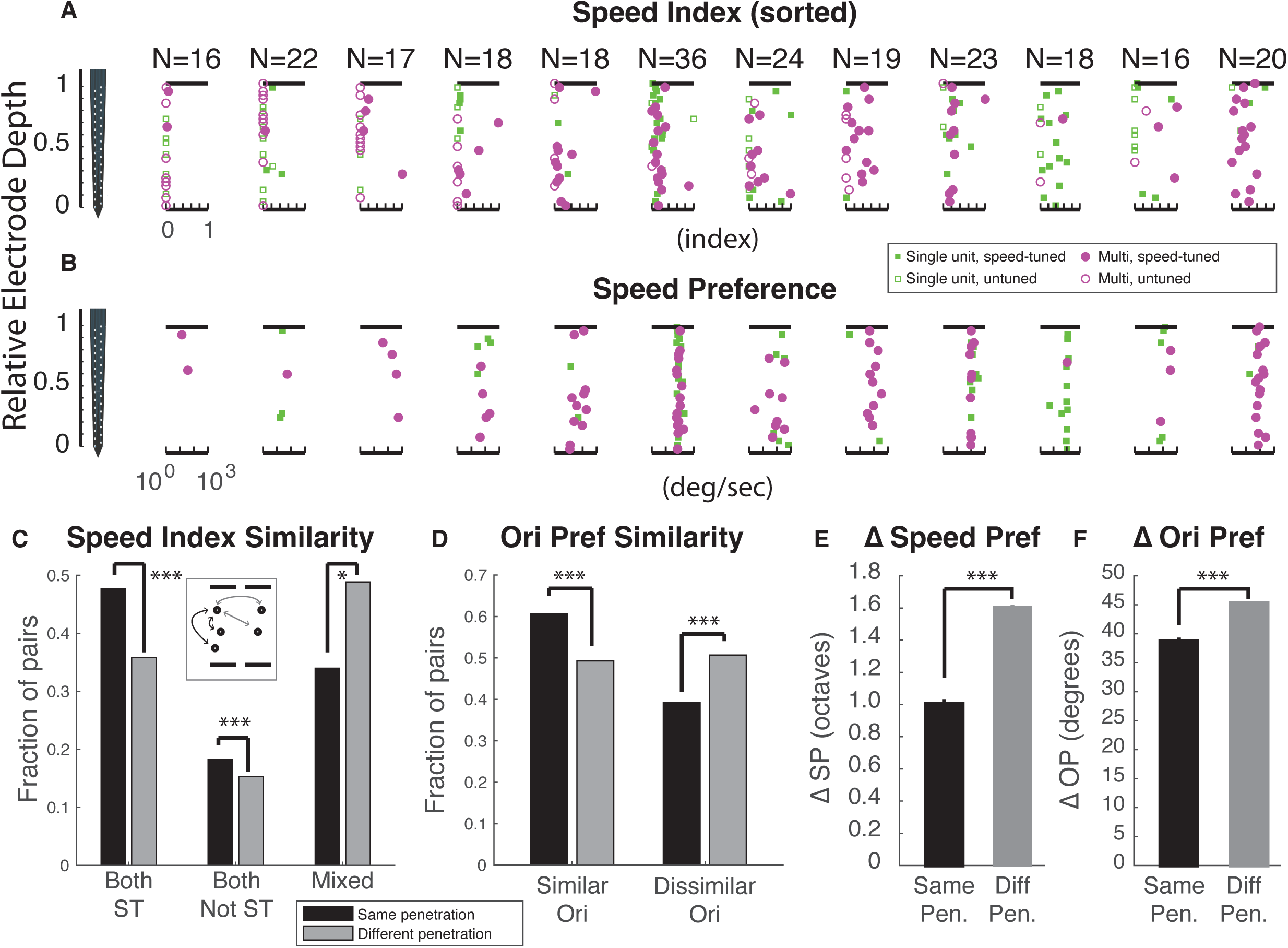
Functional architecture for speed tuning. **A)** Neuronal response properties were plotted on an index/preference (X-axis) versus relative depth along our electrode (Y-axis) to investigate potential functional architecture. Each column represents an individual penetration (n = 12 shown, n = 20 total). Single units (green squares) and multi-units (magenta circles) are displayed. Penetrations were sorted by mean speed index value. In the leftmost penetration, no cells exhibited strong speed-tuning. In the right most penetration, most cells exhibit strong speed tuning. These observations suggest that lack of speed tuning is clustered and that speed-tuning is clustered. **B)** Same, for speed preference (preferred temporal frequency divided by preferred spatial frequency) for significantly speed-tuned cells only. Columns show the same penetrations as in **A)**. As one moves left to right, the fraction of significantly speed-tuned cells increases. Most columns show tight clustering of speed preference. **C)** Speed index (SI) clustering analysis. Pairs of cells (inset) were examined either within penetrations (black) or across penetrations (gray) to determine if a) both were speed-tuned, b) neither were speed-tuned, or c) one was speed-tuned and the other wasn’t. Cell pairs where both cells were speed tuned were overrepresented within actual electrode penetrations compared to pairs taken from different penetrations (shuffle test, P<0.0001). In addition, cell pairs where neither cell exhibited significant speed tuning were also overrepresented in the actual penetrations (P<0.0001). Cell pairs that were mixed were significantly underrepresented in the actual penetrations (P<0.016). **(D)** Orientation preference clustering: Cell pairs from the same penetration were more likely to have orientation preferences within 45° (similar) than those across penetrations (shuffle test, P<0.0001), and cell pairs were likely to have dissimilar preferences (45° or greater) across different penetrations (shuffle test, P<0.0001). **(E)** Average difference in speed preference in octaves for cell pairs within penetrations as compared to cell pairs across different penetrations (shuffle test, P<0.0001). **F)** Same for orientation preference, except difference in preference is calculated in circular space.

To quantitatively probe whether speed tuned cells were clustered, we performed an analysis of the similarity of cell pairs, where the members of each pair were selected from either the same penetration (black) or different (gray) penetrations (**Figure 3C**). We compared the speed index from both cells to classify the pair as a) both significantly speed-tuned, b) both not significantly speed-tuned, or c) mixed (one cell speed-tuned while the other was not). If there were clustering, we would expect to see pairs that were “like” (both speed-tuned or both not) at a higher rate among pairs taken from the same penetration than different penetration. Consistent with this idea, the actual data exhibited significantly more clustering than in a surrogate set of simulations where the data was shuffled among penetrations (shuffle test, p < 1e-4 for both speed-tuned, p<1e-4 for both not speed-tuned, and p<0.016 for mixed) supporting the presence of speed clusters. In particular, mixed cell pairs were highly underrepresented in the actual data compared to the shuffled data, consistent with columnar organization.

To compare the clustering of speed index values against a property that is well known to exhibit a column organization we performed an analogous clustering analysis comparing the difference in orientation preference between adjacent cell pairs (black) that exhibited orientation angle preference that were similar (orientation angle preferences within 45°) or different (orientation angle preferences ≥ 45°) (**Figure 3D**). We found that adjacent cells within the same recording exhibited significantly similar tuning in orientation preference compared to those cells paired to shuffled cell pairs, showing that this style of analysis is capable of finding the known organization in orientation (shuffle tests, p<0.0001).

Lastly, we probed for clustered organization by assessing the change in speed preference of speed-tuned cells within a penetration (**Figure 3E**). We calculated the average difference (in octaves) of speed preferences for cell pairs within a penetration or across penetrations and found that cell pairs within a penetration exhibited significantly similar speed preferences as compared to cell pairs across penetrations (shuffle test, p<0.0001). As a control measurement, we also computed the average change in orientation preference within and across penetrations; as expected, cell pairs within a penetration exhibited more similar tuning than those across penetrations (shuffle test, p<0.0001).

### Laminar organization of speed index values

Our findings highlight the existence of speed-selective clusters in V1 and argues for a functional organization for speed tuning across the cortical surface. However, it remains unclear whether such selectivity also varies across cortical layers.

To assess whether speed tuning differs by laminar, we assigned each recorded cell to a standardized cortical depth. This was achieved by generating current source density (CSD) plots in response to a full-field stimulus (100 ms ON, 200 ms OFF; 50 repetitions) presented during the recording sessions. Layer 4 was identified based on characteristic current sinks following visual stimulation (Stoelzel et al., 2008) (**Figure 4A, 4B**). Using CSD landmarks, we mapped each neuron’s depth to a normalized cortical axis (**Figure 4C, 4D**), dividing the cortex into three regions: supragranular layers (L2/3), granular layer (L4), and infragranular layers (L5/6), indicated by dashed lines.

**Figure 4.**
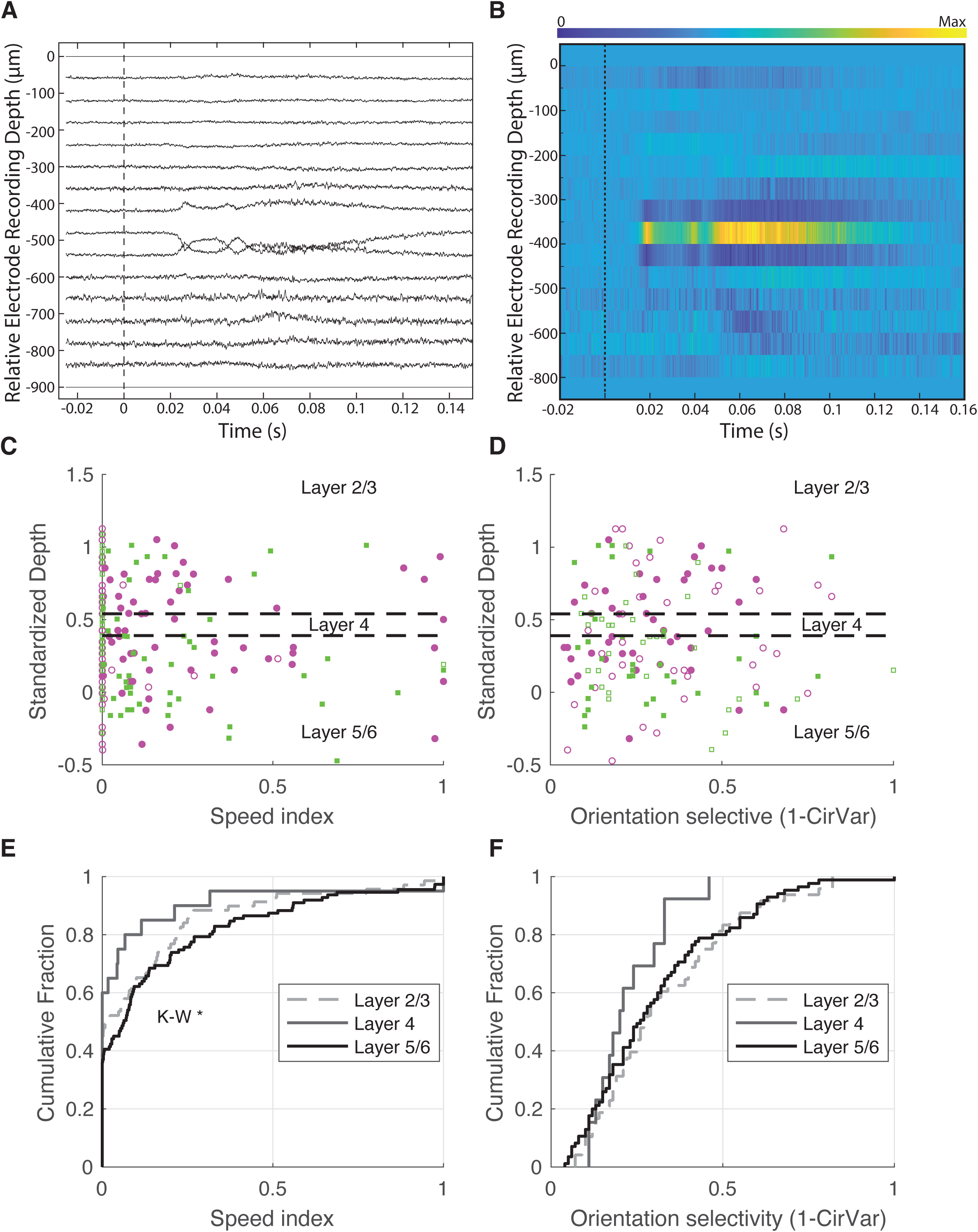
Laminar analysis of speed tuning within primary visual cortex shows modest impact of layer. **(A-B)** Identification of cortical layers using current source density (CSD) plots during full-screen blinking stimulus presentations. Time 0 indicates a full-screen black-to-white transition. The presence of a major current sink tightly locked to visual stimulus onset allowed for Layer 4 to be identified, aiding in the projection of cell depths onto standardized cortical layers. This was used to investigate how speed index values **(C)** and orientation selectivity index values (1-CirVar) **(D)** were organized according to cortical layers. **(E)** Comparison of speed index values across cortical layers (L2/3, L4, L5/6). A Kruskal-Wallis test showed evidence of a significance difference across all 3 groups (p<0.0083), although post-hoc tests showed no significant differences in speed tuning in pairwise comparisons layers (L2/3 against L4, Kruskal-Wallis Test, p = 0.3354, L2/3 against L5/6, Kruskal-Wallis Test, p = 0.3814, L4 against L5/6, Kruskal-Wallis Test, p = 0.1262). **(F)** Orientation tuning strength, measured as 1 minus circular variance. No significant differences were observed across layers (Kruskal-Wallis test, p<0.2590).

We first examined whether speed tuning varied across these layers. **Figure 4E** shows the distribution of speed index values across L2/3 (dashed), L4 (gray), and L5/6 (black). A Kruskal-Wallis Test across all layers indicating significant differences across the layers (P<0.0083). Despite the hint of a trend for L4 to exhibit less speed tuning, all of our post-hoc tests among the layers failed to reach significance, either when comparing L2/3 against L4 (K-W Test, p = 0.3354), L2/3 against L5/6 (K-W Test, p = 0.3814), or L4 against L5/6 (K-W Test, p = 0.1262).

We found no evidence of differences in orientation selectivity across the layers (K-W test, p=0.2590, **Figure 4F**).

### Evidence of speed tuning maps in cats

The presence of speed-tuned cluster in ferret V1 led us to ask whether such functional organization might extend across the cortical surface in the form of speed-tuning maps. To investigate this possibility, we reanalyzed intrinsic signal imaging data from previous studied in cat V1 (Ribot et al., 2016; Sauvage et al., 2017; Cosnier-Horeau et al., 2024). These datasets include responses to drifting gratings with co-varied spatial frequency, temporal frequency, and direction, making them suitable for pixelwise speed tuning analysis.

We fitted the intrinsic signal imaging response of each pixel with the same speed tuning function as for electrophysiology (Priebe et al., 2006). Consistent with our results in the ferret, the resulting maps revealed distinct hot spots of high speed index and cold spots of low speed index for 3 animals (**Figure 5A,D,G**), which were repeated across the cortical surface. We plotted speed preference across the cortical surface by filtering the maps for pixels that indicated significant speed tuning by a nested F test (see methods), and plotting the speed preference for the significant pixels (**Figure 5B, E, H**). The hot spots typically showed preference for relatively slow speeds (2-10 degrees per second) (**Figure 5C, F, I**).

**Figure 5.**
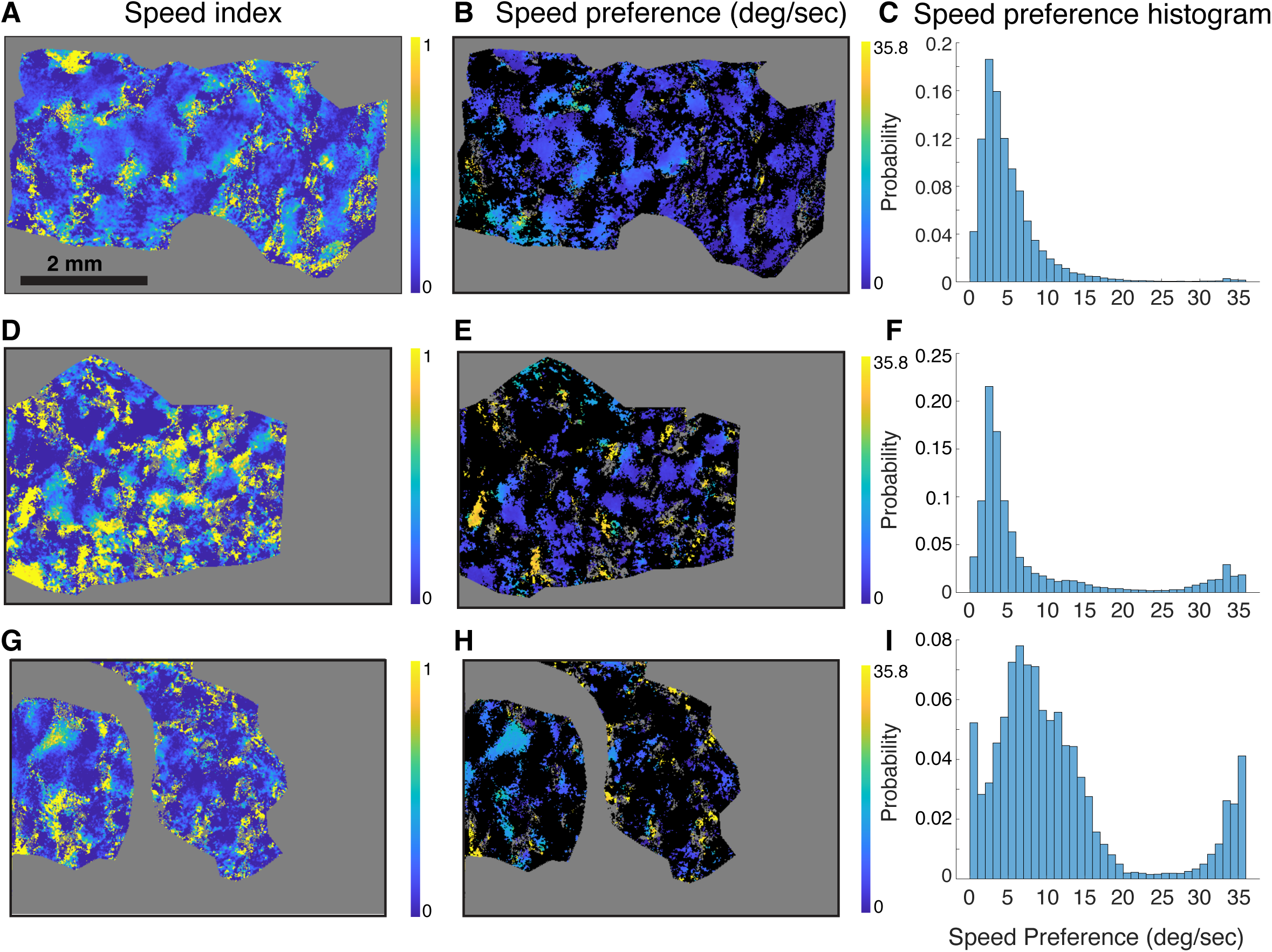
Speed tuning hot spots in cat primary visual cortex. **(A)** False-color map of speed tuning index from pixel-by-pixel fits of the spatial and temporal frequency data at the preferred direction according to the function from Priebe et al. (2006). Pixels with fits that have low r^2^ are shown in gray, as are pixels outside the region of interest. Repeating hot spots with high speed-tuning index values were observed across the cortical surface, along with large regions with low speed-tuning index values. **(B)** Speed preference for pixels that exhibit speed tuning as determined by a nested F test. **(C)** Histogram of speed preference values for pixels with significant speed tuning. **D-I**) Same, for two additional experimental animals. All cats exhibit an apparent periodic structure. Most commonly, speed preference values ranged from 2 – 10 degrees per second.

We quantified the degree of periodicity by analyzing a binary image of the pixels that indicated significant speed tuning by the nested F test analysis (**Figure 6A**). We computed an autocorrelation image (**Figure 6C, G, K**) that showed evidence of repeating regions of high and low correlation. The hot zones (**Figure 6A**) exhibited substantial power at spatial frequencies between 0.5 – 2 cycles per millimeter (**Figure 6D, H, L**). For comparison, we also quantified structure of direction maps by analyzing a binary image of pixels where direction preference values were found to be between −30° and +30° (**Figure 6B**). As expected, autocorrelation coefficient maps showed strong periodicity (**Figure 6E,I,M**) and had spatial frequency peaks in a similar range as those found for speed tuning (**Figure 6F,J,N**). Therefore, speed tuning is found in periodic hot spots across the cortical surface in cat V1.

**Figure 6.**
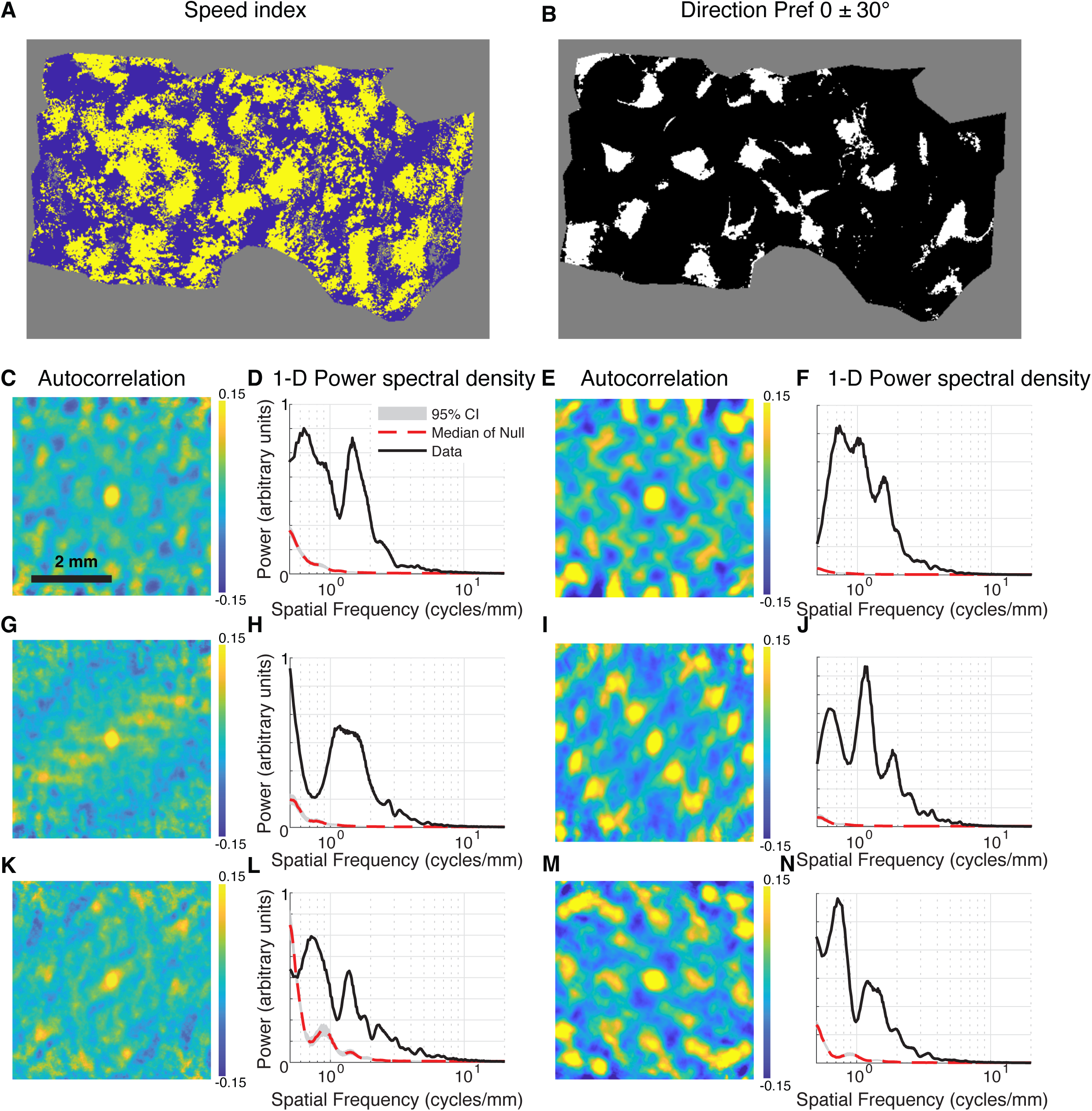
Quantitative evidence of periodic speed tuning in cat V1. **(A)** Thresholded image of regions that exhibit significant speed tuning by a nested F test (yellow) and regions that do not (blue) for one subject. Pixels out of region of interest or that did not exceed a goodness of fit of R^2^>0.2 are gray. **(B)** Thresholded image of regions that have a direction preference value that is within 30° of 0 (white) or not (black). Same subject as in (A). **(C)** Autocorrelation coefficient map of the image in (A) showing repeating regions of high and low correlation. **(D)** Power spectral density vs. spatial frequency for the image in (A). To evaluate significance, the power spectral density for 1000 surrogate images where the locations of the positive pixels are shuffled are shown in gray (range from 2.5-to-97.5 percentile) and the median by the red dashed line. The low frequency signal in the surrogate data comes from the rectangular viewing window and the shape of the region of interest. **(E)** Autocorrelation for the image in (B), also showing clear periodicity. **(F)** Power spectral density vs. spatial frequency for the image in (B). **G-N** are same for additional subjects. Regions of significant speed tuning and direction maps exhibit significant spectral power in the range of 0.5 – 2 cycles per millimeter.

### No apparent interactions among map parameters

Having established that speed index values are organized periodically in cat visual cortex, we went on to explore whether there were any obvious interactions between speed-tuned regions and the traditional parameters of orientation or direction index values or spatial frequency or temporal frequency preferences (Blasdel and Salama, 1986; Bonhoeffer and Grinvald, 1991). In **Supplementary Figure 1**, we show speed tuning maps juxtaposed to these well-studied measures. No relationships among these variables were immediate apparent. To study these relationships quantitatively, we plotted correlation density histograms among many parameters for all animals in **Supplementary Figure 2**. While occasional relationships exhibited statistically significant correlations in one animal or another, there were no strong systematic relationships that held across all animals.

## Discussion

In this study, we provided new evidence that functional architecture for speed-tuned cells exist within the primary visual cortex (V1) of ferrets, challenging the traditional view that speed processing is primarily restricted to higher-order visual areas such as MT/V5 (primate) or PMLS (cat) or PSS (ferret). Our findings showed that both simple and complex cells in V1 can exhibit speed tuning, although it is more prevalent among complex cells. We demonstrated that speed-tuned cells are spatially organized into discrete clusters or “hot spots,” but we found no evidence for a specific laminar organization. Analysis of intrinsic signal imaging results from the cat showed evidence for periodic repetition of speed-tuned hot spots and spots untuned for speed center, indicating a relationship between the zones tuned for speed and orientation maps. These findings expand our understanding of V1’s role in motion processing and suggest that functional architecture for speed processing is found earlier in the visual pathway than previously thought.

### Speed Tuning in V1

In the primate, the middle temporal area (Allman and Kaas, 1971; Born and Bradley, 2005; Zeki, 2015) contains neurons tuned for direction (Dubner and Zeki, 1971; Baker et al., 1981; Van Essen et al., 1981; Maunsell and Van Essen, 1983; Felleman and Kaas, 1984), speed (Rodman and Albright, 1987; Perrone and Thiele, 2001; Liu and Newsome, 2003, 2005; Priebe et al., 2003; Krekelberg et al., 2006), and disparity (DeAngelis et al., 1999). When direct comparisons have been made between tuning properties in MT and V1, investigators have found that V1 itself also contains neurons tuned for direction (Movshon and Newsome, 1996), speed (Orban et al., 1986b; Priebe et al., 2006), and disparity (Prince et al., 2000, 2002). Priebe et al. (2006) found that speed tuning was present in V1 complex cells, and, similarly, we find speed tuning to be more common in (but not exclusive to) complex cells. What MT appears to add beyond V1 is a selectivity to more global motion that resolves ambiguity introduced by the aperture problem (Movshon et al., 1985; Rodman and Albright, 1989) and selectivity that is more influenced by stimulation in a wide surround, enhancing differences between motion in the surround and the center. The areas PSS in the ferret and PMLS in the cat are, like MT, highly myelinated, receive direction projections from V1, and contain cells enriched for direction tuning and a sensitivity to global motion as opposed to component motion (Lempel and Nielsen, 2019, 2021). Here, our data reveals the existence of speed-tuned cells in the primary visual cortex of carnivores, with a temporal frequency tuning that varies with the spatial frequency of the stimulus. This observation adds to other reports of the existence of such cells in primate V1 (Orban et al., 1986b; Priebe et al., 2006) or neurons tuned to speed *per se* in mouse (Andermann et al., 2011; Glickfeld et al., 2014; McBride et al., 2019; Wang et al., 2021), suggesting that this is a common property of sensory tuning in V1, along with selectivity for orientation, direction, disparity, and color.

### Functional architecture of speed tuning in V1

Area MT exhibits a columnar architecture for many of its tuning properties, including direction preference, speed preference, and disparity (Fennema and Thompson, 1979; Van Essen et al., 1981; Felleman and Kaas, 1984; Orban et al., 1986b, 1986b; Rodman and Albright, 1987, 1989; Lagae et al., 1989, 1993, 1993; Shipp and Zeki, 1989; Movshon and Newsome, 1996; Orban, 1997; DeAngelis and Newsome, 1999; Britten, 2003; Liu and Newsome, 2003; Churchland et al., 2005). MT in the macaque is buried in the superior temporal sulcus, so there have not been imaging studies that have precisely mapped the organization of these cells, and instead, our knowledge of this architecture comes from electrophysiology studies that incorporated serial recordings of cells along straight electrode penetrations; however see these marmoset studies (Solomon and Rosa, 2014; Mitchell and Leopold, 2015; Kwan et al., 2021). These studies have suggested that direction maps in MT might resemble those in the cat and the ferret, as investigators observed smooth progressions of direction preference across the cortical surface with occasional jumps of 180° as electrodes cross direction fractures (Blasdel and Salama, 1986; Malonek et al., 1994). Disparity tuning is organized into zones of near, far, or smooth intermediate progression (DeAngelis et al., 1998, 1999; DeAngelis and Uka, 2003). Speed tuning is also clustered, although speed preference sometimes is constant for long stretches in MT while direction preference undergoes a smooth progression (Liu and Newsome, 2003). Thus, it is unclear if speed preference in MT is organized in a smooth map with a lower rate of change than direction preference, or if instead speed preference is merely locally clustered without an overall smooth map (Born and Bradley, 2005).

Despite the known functional organization of MT, we are unaware of any previous reports of functional organization for speed tuning in the primary visual cortex, as we show here. Our electrophysiological measurements showed a clustering of speed tuned cells that was recapitulated in a reanalysis of speed tuning in intrinsic signal imaging data from cat visual cortex. Cat visual cortex exhibited similar periodicity for speed tuning as it does for other features like direction preference, indicating that speed tuning is somehow embedded into the maps of orientation, direction, and spatial frequency that are well known. Therefore, the carnivore visual cortex exhibits functional architecture for visual space (Yu et al., 2005), orientation (Bonhoeffer and Grinvald, 1991; Chapman et al., 1996; Ohki et al. 2006), direction (Weliky et al. 1996; Ohki et al. 2005), spatial and temporal frequency (Shoham et al., 1997; Hübener et al. 1997; Yu et al., 2005; Baker and Issa 2005; Farley et al. 2007; Issa et al., 2008), disparity (Kara and Boyd, 2009), and speed (this study). It remains to be seen whether these zones reflect, in carnivores, a functional division of projection patterns to extrastriate areas that resemble the projections to the thick and thin stripes of V2 in the primate (Livingstone and Hubel, 1984, 1987, 1988; DeYoe and Van Essan, 1985; Shipp and Zeik 1985, 1989; Felleman and Van Essen, 1991; Nakamura et al., 1993; Sincich and Horton, 2005).

### Speculation about circuit mechanisms of speed tuning

What circuit mechanisms might underlie the speed tuning we observe in ferret V1? In 1998, when speed-tuning was well-known in MT but still not well-known in V1, Simoncelli and Heeger proposed a model of the V1-to-MT projection (Simoncelli and Heeger, 1998, 2001; Nishimoto and Gallant, 2011). They posited that V1 complex cells that were selective for single orientations, spatial frequencies, and temporal frequencies converged onto MT cells in a selective manner. Their input V1 cells exhibited tuning only for these properties (orientation, spatial frequency, temporal frequency) and not for speed, and their input V1 cells responded to component motion when stimulated with “plaid” gratings (mixtures of 2 drifting grating stimuli of equal contrast). The MT neurons exhibited both speed tuning and tuning to the “global” motion of these plaid grating stimuli. It is well-known in primates (Movshon and Newsome, 1996) and carnivores (Lempel and Nielsen, 2021) that V1 neurons exhibit selectivity almost exclusively for component motion, while neurons MT and PSS exhibit selectivity for global motion. The widespread presence of speed-tuned cells in V1 suggests a different progression: V1 simple cells tuned for single orientations, spatial frequencies, and temporal frequencies project to V1 complex cells (Priebe et al., 2006), creating a population of speed-tuned complex cells in V1. Then, these complex cells project to MT or PSS, where the connection strength is determined by the difference in direction preference angle (high when matched, less high for a small mismatch, and low for a big mismatch). Under this scenario, MT neurons would exhibit tuning for global motion as observed, but speed tuning would be computed earlier, in V1.

On the other hand, there is evidence that MT/PSS/PLMS maintain much of their responses without V1, but also see Collins et al., 2005 (Collins et al., 2005 p.203), raising the possibility that the flow of information might at least partially run in the opposite direction (from MT/PSS/PLMS to V1). Silencing V1 does not eliminate direction selectivity in MT (Azzopardi et al., 2003; Gross et al., 2004) or PSS (Lempel and Nielsen, 2021), and removing primary visual cortex in the cat does not eliminate motion tuning in PLMS (Guido et al., 1990, 1992). Connections from the koniocellular/ W-cell layers of LGN to MT/PSS/PLMS or via superior colliculus and pulvinar might drive some of these responses (Rodman et al., 2001; Lyon et al., 2010).

Within V1, we found that some zones of V1 exhibited speed-tuning while others did not. This raises the question of whether the connectivity rules that define these receptive fields differ between these zones, or rather connectivity rules are the same across cortex and whether a zone is speed-tuned or not is simply a matter of where the zone is located in the orientation and spatial frequency map.

Finally, we found evidence that some simple cells already expressed speed tuning, although this was less common than for complex cells. Further, speed tuning index values were empirically (but not significantly) smaller in layer 4, suggesting a smaller contribution of speed tuning there. Nevertheless, this raises the question of how some simple cells, which are thought to receive input from spot-detector LGN cells (Reid and Alonso, 1995), might achieve speed tuning. Future work will be needed to understand whether feed-forward LGN projections can impart speed tuning directly (Sincich et al., 2004).

## Acknowledgements

This work was funded by NIH EY022122 (SDV) and NIH NIGMS 1T32GM132498.

## Author contributions

V.M.S.C. and S.D.V.H. designed research; V.M.S.C. and L.Y. performed research; NLN and V.M.S.C. analyzed data; V.M.S.C. and S.D.V.H. wrote the paper.

## Figure Captions

**Supplementary Figure 1.**
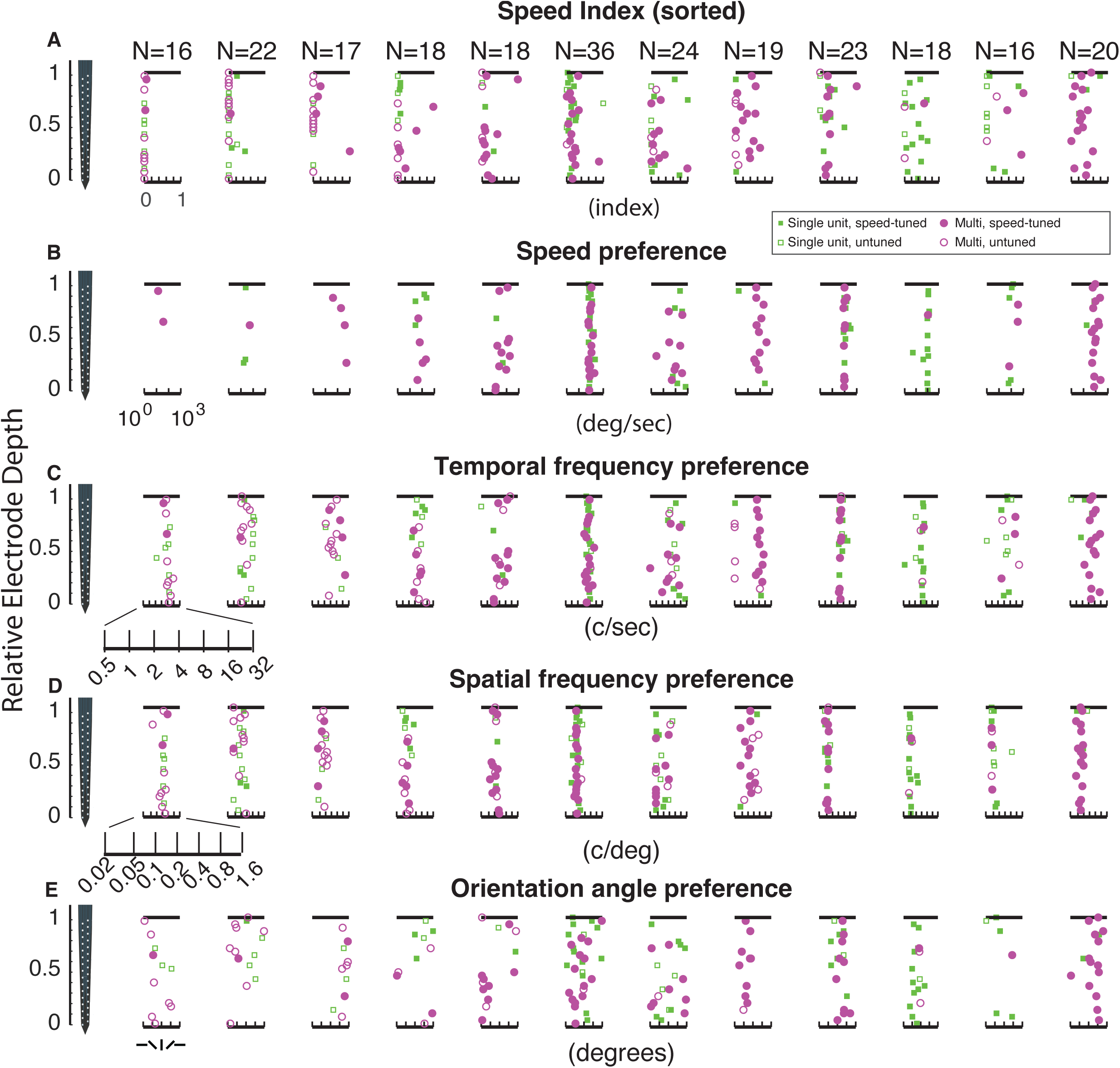
Tuning properties of cells in individual electrode penetrations. **C)** Temporal frequency preference. **D)** Spatial frequency preference. **E) O**rientation preference. All penetrations exhibit tuning properties that are more similar within the electrode penetration than across electrode penetrations.

**Supplementary Figure 2.**
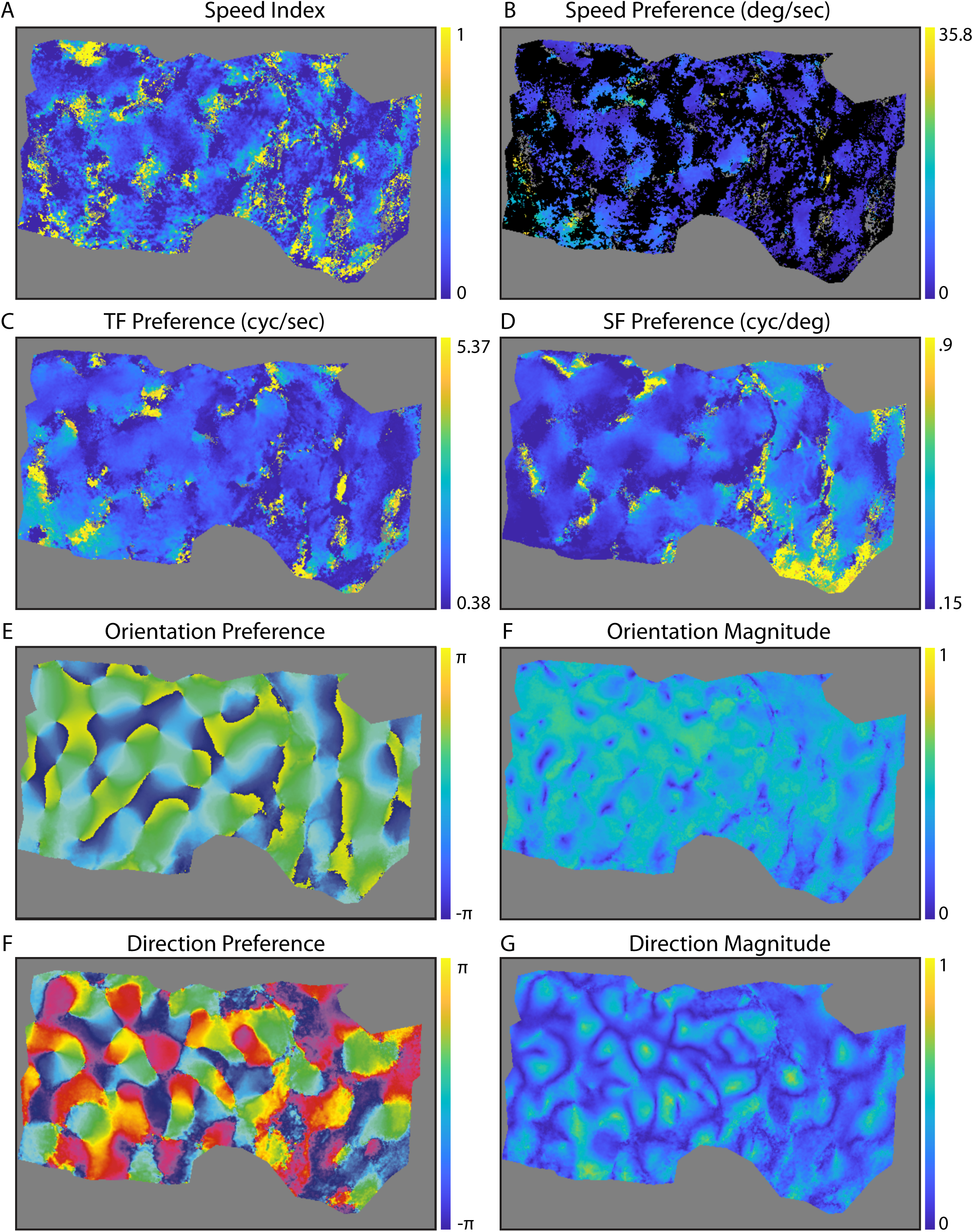
Speed tuning hot spots in cat primary visual cortex for one subject. **(A)** False-color map of speed tuning index as before. **(B)** Speed preference for pixels that exhibit significant speed tuning by the nested F test. **(C)** Preferred temporal frequency from the Priebe et al. (2006) function. Temporal frequency preference values are close to 2 Hz over much of the cortex, with some small zones of high temporal frequency preference. **(D)** Spatial frequency preference from the Priebe et al. (2006) function. There is a gradient from low (left side) to high (right side). **(E)** Orientation preference angle. **(F)** Orientation vector magnitude; orientation index values are generally high over the cortex. **(G)** Direction preference map. **(H)** Direction magnitude map. Known regions of high direction selectivity and direction fractures (regions of low direction selectivity that form lines) are observed (Weliky et al., 1996; Ohki et al., 2005).

**Supplementary Figure 3.**
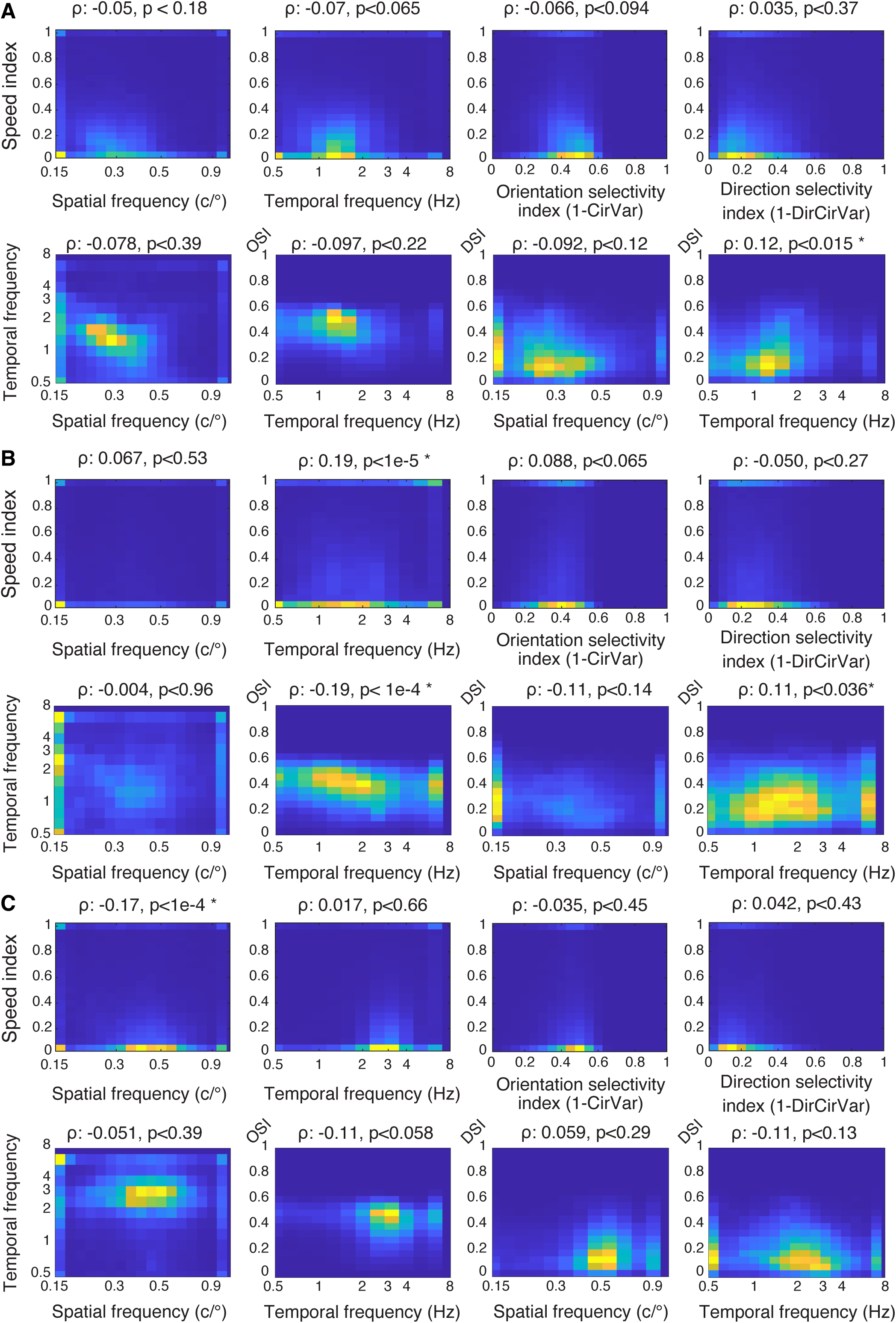
No consistent or strong correlations between speed index values and preferred spatial frequencies, preferred temporal frequencies, orientation selectivity index values, or direction selectivity index values. **A)** For one cat, densities (2-dimensional histograms) of speed index values and spatial frequency preferences, temporal frequency preferences, and 1-CirVar and 1-DirCirVar. In the second row, densities of relationships between non-speed parameters are shown. Spatial frequency, and temporal frequency are on log scales. Correlation coefficients (ρ) and p values calculated using shuffled maps to remove the autocorrelation of each parameter (see methods) are indicated. Colors indicate the number of pixels that fall into a given speed index / abscissa bin, with yellower colors indicating higher counts. **B,C)** Same, for additional animals.

## References

Allman JM, Kaas JH (1971) A representation of the visual field in the caudal third of the middle tempral gyrus of the owl monkey (Aotus trivirgatus). Brain Res 31:85–105.

Andermann ML, Kerlin AM, Roumis DK, Glickfeld LL, Reid RC (2011) Functional specialization of mouse higher visual cortical areas. Neuron 72:1025–1039.

Azzopardi P, Fallah M, Gross CG, Rodman HR (2003) Response latencies of neurons in visual areas MT and MST of monkeys with striate cortex lesions. Neuropsychologia 41:1738–1756.

Baker CL (1990) Spatial- and temporal-frequency selectivity as a basis for velocity preference in cat striate cortex neurons. Vis Neurosci 4:101–113 Available at: http://www.journals.cambridge.org/abstract_S0952523800002273 [Accessed October 25, 2022].

Baker JF, Petersen SE, Newsome WT, Allman JM (1981) Visual response properties of neurons in four extrastriate visual areas of the owl monkey (Aotus trivirgatus): a quantitative comparison of medial, dorsomedial, dorsolateral, and middle temporal areas. J Neurophysiol 45:397–416.

Baker TI, Issa NP (2005) Cortical Maps of Separable Tuning Properties Predict Population Responses to Complex Visual Stimuli. Journal of Neurophysiology 94:775–787 Available at: https://journals.physiology.org/doi/full/10.1152/jn.01093.2004 [Accessed June 3, 2021].

Blasdel GG, Salama G (1986) Voltage-sensitive dyes reveal a modular organization in monkey striate cortex. Nature 321:579–585 Available at: 10.1038/321579a0.

Bonhoeffer T, Grinvald A (1991) Iso-orientation domains in cat visual cortex are arranged in pinwheel-like patterns. Nature 353:429–431 Available at: https://www.nature.com/articles/353429a0 [Accessed December 6, 2020].

Born RT, Bradley DC (2005) Structure and function of visual area MT. Annu Rev Neurosci 28:157–189.

Brainard DH (1997) The Psychophysics Toolbox. Spat Vis 10:433–436.

Britten K (2003) The middle temporal area: Motion processing and the link to perception. The visual neurosciences.

Chapman B, Stryker MP (1993) Development of orientation selectivity in ferret visual cortex and effects of deprivation. J Neurosci 13:5251–5262.

Chapman B, Stryker MP, Bonhoeffer T (1996) Development of orientation preference maps in ferret primary visual cortex. J Neurosci 16:6443–6453.

Churchland MM, Priebe NJ, Lisberger SG (2005) Comparison of the spatial limits on direction selectivity in visual areas MT and V1. Journal of Neurophysiology 93:1235–1245.

Collins CE, Lyon DC, Kaas JH (2005) Distribution across cortical areas of neurons projecting to the superior colliculus in new world monkeys. Anat Rec A Discov Mol Cell Evol Biol 285:619–627.

Cosnier-Horeau C, Germaine H, Van Hooser SD, Choi I, Ribot J, Touboul J (2024) Latent coding of movement in primary visual cortex. bioRxiv 2024.11.11.623057

DeAngelis GC, Cumming BG, Newsome WT (1998) Cortical area MT and the perception of stereoscopic depth. Nature 394:677–680.

DeAngelis GC, Ghose GM, Ohzawa I, Freeman RD (1999) Functional Micro-Organization of Primary Visual Cortex: Receptive Field Analysis of Nearby Neurons. J Neurosci 19:4046– 4064.

DeAngelis GC, Newsome WT (1999) Organization of disparity-selective neurons in macaque area MT. Journal of Neuroscience 19:1398–1415.

DeAngelis GC, Uka T (2003) Coding of horizontal disparity and velocity by MT neurons in the alert macaque. Journal of neurophysiology 89:1094–1111.

DeYoe, E. A., & Van Essen, D. C. (1985). Segregation of efferent connections and receptive field properties in visual area V2 of the macaque. Nature, 317(6032), 58–61.

DiCarlo JJ, Lane JW, Hsiao SS, Johnson KO (1996) Marking microelectrode penetrations with fluorescent dyes. Journal of Neuroscience Methods 64:75–81 Available at: https://www.sciencedirect.com/science/article/pii/0165027095001131 [Accessed September 3, 2024].

Dubner R, Zeki SM (1971) Response properties and receptive fields of cells in an anatomically defined region of the superior temporal sulcus in the monkey. Brain Res 35:528–532.

Farley, B. J., Yu, H., Jin, D. Z., & Sur, M. (2007). Alteration of visual input results in a coordinated reorganization of multiple visual cortex maps. The Journal of neuroscience 27(38), 10299–10310. 10.1523/JNEUROSCI.2257-07.2007

Felleman DJ, Kaas JH (1984) Receptive-field properties of neurons in middle temporal visual area (MT) of owl monkeys. J Neurophysiol 52:488–513.

Felleman, D. J., & Van Essen, D. C. (1991). Distributed hierarchical processing in the primate cerebral cortex. Cerebral Cortex, 1(1), 1–47.

Fennema CL, Thompson WB (1979) Velocity determination in scenes containing several moving objects. Computer graphics and image processing 9:301–315.

García Murillo, D., Zhao, Y., Rogovin, O. S., Zhang, K., Hu, A. W., Kim, M. R., Chen, S., Wang, Z., Keeley, Z. C., Shin, D. I., Suárez Casanova, V. M., Zhu, Y., Martin, L., Papaemmanouil, O., & Van Hooser, S. D. (2022). NDI: A Platform-Independent Data Interface and Database for Neuroscience Physiology and Imaging Experiments. eNeuro, 9(1), ENEURO.0073-21.2022. 10.1523/ENEURO.0073-21.2022

Gattass, R., Sousa, A. P., Mishkin, M., & Ungerleider, L. G. (1997). Cortical projections of area V2 in the macaque. Cerebral Cortex, 7(2), 110–129.

Gilardi C, Kalebic N (2021) The Ferret as a Model System for Neocortex Development and Evolution. Front Cell Dev Biol 9 Available at: https://www.frontiersin.org/articles/10.3389/fcell.2021.661759/full [Accessed June 2, 2021].

Glickfeld LL, Reid RC, Andermann ML (2014) A mouse model of higher visual cortical function. Curr Opin Neurobiol 24:28–33.

Gross CG, Moore T, Rodman HR (2004) Visually guided behavior after V1 lesions in young and adult monkeys and its relation to blindsight in humans. Prog Brain Res 144:279–294.

Guido W, Spear PD, Tong L (1990) Functional compensation in the lateral suprasylvian visual area following bilateral visual cortex damage in kittens. Exp Brain Res 83:219–224.

Guido W, Spear PD, Tong L (1992) How complete is physiological compensation in extrastriate cortex after visual cortex damage in kittens? Exp Brain Res 91:455–466.

Hawken MJ, Parker AJ, Lund JS (1988) Laminar organization and contrast sensitivity of direction-selective cells in the striate cortex of the Old World monkey. J Neurosci 8:3541–3548 Available at: https://www-jneurosci-org.eu1.proxy.openathens.net/content/8/10/3541 [Accessed June 2, 2021].

Heimel JA, Van Hooser SD, Nelson SB (2005) Laminar organization of response properties in primary visual cortex of the gray squirrel (Sciurus carolinensis). J Neurophysiol 94:3538–3554.

Hubel DH, Wiesel TN (1959) Receptive fields of single neurones in the cat’s striate cortex. The Journal of Physiology 148:574–591 Available at: https://onlinelibrary.wiley.com/doi/abs/10.1113/jphysiol.1959.sp006308 [Accessed August 30, 2024].

Hubel DH, Wiesel TN (1962) Receptive fields, binocular interaction and functional architecture in the cat’s visual cortex. The Journal of Physiology 160:106–154 Available at: https://onlinelibrary.wiley.com/doi/abs/10.1113/jphysiol.1962.sp006837 [Accessed August 30, 2024].

Hubel DH, Wiesel TN (1963) Shape and arrangement of columns in cat’s striate cortex. The Journal of Physiology 165:559–568 Available at: https://onlinelibrary.wiley.com/doi/abs/10.1113/jphysiol.1963.sp007079 [Accessed September 13, 2024].

Hubel DH, Wiesel TN (1965) RECEPTIVE FIELDS AND FUNCTIONAL ARCHITECTURE IN TWO NONSTRIATE VISUAL AREAS (18 AND 19) OF THE CAT. J Neurophysiol 28:229– 289.

Hubel DH, Wiesel TN (1968) Receptive fields and functional architecture of monkey striate cortex. The Journal of Physiology 195:215–243 Available at: https://physoc.onlinelibrary.wiley.com/doi/abs/10.1113/jphysiol.1968.sp008455 [Accessed June 2, 2021].

Hubel DH, Wiesel TN (1997) Ferrier lecture - Functional architecture of macaque monkey visual cortex. Proceedings of the Royal Society of London Series B Biological Sciences 198:1– 59 Available at: https://royalsocietypublishing.org/doi/10.1098/rspb.1977.0085 [Accessed August 30, 2024].

Hubel DH, Wiesel TN (1998) Early Exploration of the Visual Cortex. Neuron 20:401–412 Available at: https://www.sciencedirect.com/science/article/pii/S0896627300809848 [Accessed August 30, 2024].

Hübener M, Shoham D, Grinvald A, Bonhoeffer T (1997) Spatial relationships among three columnar systems in cat area 17. J Neurosci 17:9270–9284.

Issa NP, Rosenberg A, Husson TR (2008) Models and measurements of functional maps in V1. J Neurophysiol 99:2745–2754.

Issa NP, Trachtenberg JT, Chapman B, Zahs KR, Stryker MP (1999) The critical period for ocular dominance plasticity in the Ferret’s visual cortex. J Neurosci 19:6965–6978.

Jun JJ, Mitelut C, Lai C, Gratiy SL, Anastassiou CA, Harris TD (2017) Real-time spike sorting platform for high-density extracellular probes with ground-truth validation and drift correction.: 101030 Available at: https://www.biorxiv.org/content/10.1101/101030v2 [Accessed August 30, 2024].

Kalatsky VA, Stryker MP (2003) New paradigm for optical imaging: temporally encoded maps of intrinsic signal. Neuron 38:529–545.

Kara, P., & Boyd, J. D. (2009). A micro-architecture for binocular disparity and ocular dominance in visual cortex. Nature, 458(7238), 627–631. 10.1038/nature07721

Katz LC, Crowley JC (2002) Development of cortical circuits: lessons from ocular dominance columns. Nat Rev Neurosci 3:34–42.

Krekelberg B, van Wezel RJA, Albright TD (2006) Interactions between speed and contrast tuning in the middle temporal area: implications for the neural code for speed. J Neurosci 26:8988–8998.

Kwan WC, Chang C-K, Yu H-H, Mundinano IC, Fox DM, Homman-Ludiye J, Bourne JA (2021) Visual Cortical Area MT Is Required for Development of the Dorsal Stream and Associated Visuomotor Behaviors. J Neurosci 41:8197–8209 Available at: https://www.jneurosci.org/content/41/39/8197 [Accessed September 18, 2024].

Lagae L, Gulyas B, Raiguel S, Orban G (1989) Laminar analysis of motion information processing in macaque V5. Brain research 496:361–367.

Lagae L, Raiguel S, Orban G (1993) Speed and direction selectivity of macaque middle temporal neurons. Journal of neurophysiology 69:19–39.

Lempel AA, Nielsen KJ (2019) Ferrets as a Model for Higher-Level Visual Motion Processing. Curr Biol 29:179–191.e5.

Lempel AA, Nielsen KJ (2021) Development of visual motion integration involves coordination of multiple cortical stages Baker CI, Pasternak T, Goris R, eds. eLife 10:e59798 Available at: 10.7554/eLife.59798 [Accessed September 3, 2024].

Liu J, Newsome WT (2003) Functional organization of speed tuned neurons in visual area MT. J Neurophysiol 89:246–256.

Liu J, Newsome WT (2005) Correlation between speed perception and neural activity in the middle temporal visual area. J Neurosci 25:711–722.

Liu L (2019) Painting Neuropixels probes and other silicon probes for electrophysiological recordings. Available at: https://www.protocols.io/view/painting-neuropixels-probes-and-other-silicon-prob-wxqffmw [Accessed April 5, 2023].

Livingstone, M. S., & Hubel, D. H. (1984). Anatomy and physiology of a color system in the primate visual cortex. Journal of Neuroscience, 4(1), 309–356.

Livingstone, M. S., & Hubel, D. H. (1988). Segregation of form, color, movement, and depth: anatomy, physiology, and perception. Science, 240(4853), 740–749.

Livingstone, M. S., & Hubel, D. H. (1987). Connections between layer 4B of area 17 and the thick cytochrome oxidase stripes of area 18 in the squirrel monkey. Journal of Neuroscience, 7(11), 3371–3377.

Lyon DC, Nassi JJ, Callaway EM (2010) A disynaptic relay from superior colliculus to dorsal stream visual cortex in macaque monkey. Neuron 65:270–279.

Maier A, Adams GK, Aura C, Leopold DA (2010) Distinct Superficial and Deep Laminar Domains of Activity in the Visual Cortex during Rest and Stimulation. Front Syst Neurosci 4 Available at: https://www.frontiersin.org/journals/systems-neuroscience/articles/10.3389/fnsys.2010.00031/full [Accessed September 18, 2024].

Maier A, Aura CJ, Leopold DA (2011) Infragranular Sources of Sustained Local Field Potential Responses in Macaque Primary Visual Cortex. J Neurosci 31:1971–1980 Available at: https://www.jneurosci.org/content/31/6/1971 [Accessed September 18, 2024].

Malonek D, Tootell R, Grinvald A (1994) Optical imaging reveals the functional architecture of neurons processing shape and motion in owl monkey area MT. Proceedings of the Royal Society of London Series B: Biological Sciences 258:109–119.

Maunsell JH, Van Essen DC (1983) Functional properties of neurons in middle temporal visual area of the macaque monkey. I. Selectivity for stimulus direction, speed, and orientation. Journal of Neurophysiology 49:1127–1147 Available at: https://journals.physiology.org/doi/abs/10.1152/jn.1983.49.5.1127 [Accessed August 30, 2024].

Mazurek M, Kager M, Van Hooser SD (2014) Robust quantification of orientation selectivity and direction selectivity. Front Neural Circuits 8:92.

McBride EG, Lee S-YJ, Callaway EM (2019) Local and Global Influences of Visual Spatial Selection and Locomotion in Mouse Primary Visual Cortex. Curr Biol 29:1592–1605.e5.

Milleret C (2017) Traitement physiologique de l’information visuelle. In: Déficiences Visuelles; Rapport de la Société française d’ophtalmologie (Société française d’ophtalmologie, ed). Issy-les-Moulineaux: Elsevier Masson.

Mitchell JF, Leopold DA (2015) The marmoset monkey as a model for visual neuroscience. Neurosci Res 93:20–46.

Moore BD (2006) Speed Selectivity in V1: A Complex Affair. J Neurosci 26:7543–7544 Available at: https://www.jneurosci.org/content/26/29/7543 [Accessed May 25, 2021].

Moore BD, Alitto HJ, Usrey WM (2005) Orientation Tuning, But Not Direction Selectivity, Is Invariant to Temporal Frequency in Primary Visual Cortex. Journal of Neurophysiology 94:1336–1345 Available at: https://journals.physiology.org/doi/full/10.1152/jn.01224.2004 [Accessed May 17, 2021].

Movshon JA, Adelson EH, Gizzi MS, Newsome WT (1985) The Analysis of Moving Visual Patterns. In: Pattern Recognition Mechanisms (Chagas C, Gattass R, Gross C, eds), pp 117–151 Experimental Brain Research Supplementum. Berlin, Heidelberg: Springer Berlin Heidelberg. Available at: http://link.springer.com/10.1007/978-3-662-09224-8_7 [Accessed September 3, 2024].

Movshon JA, Newsome WT (1996) Visual Response Properties of Striate Cortical Neurons Projecting to Area MT in Macaque Monkeys. J Neurosci 16:7733–7741 Available at: https://www.jneurosci.org/content/16/23/7733 [Accessed August 30, 2024].

Movshon JA, Thompson ID, Tolhurst DJ (1978a) Spatial and temporal contrast sensitivity of neurones in areas 17 and 18 of the cat’s visual cortex. J Physiol 283:101–120.

Movshon JA, Thompson ID, Tolhurst DJ (1978b) Receptive field organization of complex cells in the cat’s striate cortex. J Physiol 283:79–99.

Movshon JA, Thompson ID, Tolhurst DJ (1978c) Spatial summation in the receptive fields of simple cells in the cat’s striate cortex. J Physiol 283:53–77.

Nakamura, H., Gattass, R., Desimone, R., & Ungerleider, L. G. (1993). The modular organization of projections from areas V1 and V2 to areas V4 and TEO in macaques. Journal of Neuroscience, 13(9), 3681–3691.

Nishimoto S, Gallant JL (2011) A Three-Dimensional Spatiotemporal Receptive Field Model Explains Responses of Area MT Neurons to Naturalistic Movies. J Neurosci 31:14551– 14564 Available at: https://www.jneurosci.org/content/31/41/14551 [Accessed August 30, 2024].

Ohki K, Chung S, Ch’ng YH, Kara P, Reid RC (2005) Functional imaging with cellular resolution reveals precise micro-architecture in visual cortex. Nature 433:597–603.

Ohki, K., Chung, S., Kara, P., Hübener, M., Bonhoeffer, T., & Reid, R. C. (2006). Highly ordered arrangement of single neurons in orientation pinwheels. Nature, 442(7105), 925–928. 10.1038/nature05019.

Orban GA (1997) Visual processing in macaque area MT/V5 and its satellites (MSTd and MSTv). In: Extrastriate cortex in primates, pp 359–434. Springer.

Orban GA, Kennedy H, Bullier J (1986a) Velocity sensitivity and direction selectivity of neurons in areas V1 and V2 of the monkey: influence of eccentricity. Journal of Neurophysiology 56:462–480 Available at: https://journals.physiology.org/doi/abs/10.1152/jn.1986.56.2.462 [Accessed September 6, 2024].

Orban GA, Kennedy H, Bullier J (1986b) Velocity sensitivity and direction selectivity of neurons in areas V1 and V2 of the monkey: influence of eccentricity. Journal of Neurophysiology 56:462–480 Available at: https://journals.physiology.org/doi/abs/10.1152/jn.1986.56.2.462 [Accessed August 30, 2024].

Pelli DG (1997) The VideoToolbox software for visual psychophysics: transforming numbers into movies. Spat Vis 10:437–442.

Perrone JA (2006) A single mechanism can explain the speed tuning properties of MT and V1 complex neurons. J Neurosci 26:11987–11991.

Perrone JA, Thiele A (2001) Speed skills: measuring the visual speed analyzing properties of primate MT neurons. Nat Neurosci 4:526–532.

Price NSC, Crowder NA, Hietanen MA, Ibbotson MR (2006) Neurons in V1, V2, and PMLS of cat cortex are speed tuned but not acceleration tuned: the influence of motion adaptation. J Neurophysiol 95:660–673.

Priebe NJ, Cassanello CR, Lisberger SG (2003) The Neural Representation of Speed in Macaque Area MT/V5. J Neurosci 23:5650–5661 Available at: https://www.jneurosci.org/content/23/13/5650 [Accessed May 25, 2021].

Priebe NJ, Lisberger SG, Movshon JA (2006) Tuning for spatiotemporal frequency and speed in directionally selective neurons of macaque striate cortex. J Neurosci 26:2941–2950.

Prince S j. d., Cumming BG, Parker AJ (2002) Range and Mechanism of Encoding of Horizontal Disparity in Macaque V1. Journal of Neurophysiology 87:209–221 Available at: https://journals.physiology.org/doi/full/10.1152/jn.00466.2000 [Accessed September 3, 2024].

Prince SJD, Pointon AD, Cumming BG, Parker AJ (2000) The Precision of Single Neuron Responses in Cortical Area V1 during Stereoscopic Depth Judgments. J Neurosci 20:3387–3400 Available at: https://www.jneurosci.org/content/20/9/3387 [Accessed September 3, 2024].

Radtke-Schuller S (2018) Cyto- and Myeloarchitectural Brain Atlas of the Ferret (Mustela putorius) in MRI Aided Stereotaxic Coordinates. Cham: Springer International Publishing. Available at: http://link.springer.com/10.1007/978-3-319-76626-3 [Accessed August 30, 2024].

Rao SC, Toth LJ, Sur M (1997) Optically imaged maps of orientation preference in primary visual cortex of cats and ferrets. J Comp Neurol 387:358–370.

Reid RC, Alonso JM (1995) Specificity of monosynaptic connections from thalamus to visual cortex. Nature 378:281–284.

Ribot J, Romagnoni A, Milleret C, Bennequin D, Touboul J (2016) Pinwheel-dipole configuration in cat early visual cortex. Neuroimage 128:63–73.

Ringach DL, Shapley RM, Hawken MJ (2002) Orientation Selectivity in Macaque V1: Diversity and Laminar Dependence. J Neurosci 22:5639–5651 Available at: https://www.jneurosci.org/content/22/13/5639 [Accessed December 13, 2023].

Ritter NJ, Anderson NM, Hooser SDV (2017) Visual Stimulus Speed Does Not Influence the Rapid Emergence of Direction Selectivity in Ferret Visual Cortex. J Neurosci 37:1557– 1567 Available at: https://www.jneurosci.org/content/37/6/1557 [Accessed January 17, 2023].

Rodman HR, Albright TD (1987) Coding of visual stimulus velocity in area MT of the macaque. Vision Res 27:2035–2048.

Rodman HR, Albright TD (1989) Single-unit analysis of pattern-motion selective properties in the middle temporal visual area (MT). Exp Brain Res 75:53–64.

Rodman HR, Sorenson KM, Shim AJ, Hexter DP (2001) Calbindin immunoreactivity in the geniculo-extrastriate system of the macaque: implications for heterogeneity in the koniocellular pathway and recovery from cortical damage. J Comp Neurol 431:168–181.

Sauvage A, Hubert G, Touboul J, Ribot J (2017) The hemodynamic signal as a first-order low-pass temporal filter: Evidence and implications for neuroimaging studies. Neuroimage 155:394– 405.

Shipp, S., & Zeki, S. (1985). Segregation of pathways leading from area V2 to areas V4 and V5 of macaque monkey visual cortex. Nature, 315(6017), 322–325.

Shipp S, Zeki S (1989) The organization of connections between areas V5 and V1 in macaque monkey visual cortex. European Journal of Neuroscience 1:309–332.

Shoham, D., Hübener, M., Schulze, S., Grinvald, A., & Bonhoeffer, T. (1997). Spatio-temporal frequency domains and their relation to cytochrome oxidase staining in cat visual cortex. Nature, 385(6616), 529–533. 10.1038/385529a0

Simoncelli EP, Heeger DJ (1998) A model of neuronal responses in visual area MT. Vision Res 38:743– 761.

Simoncelli EP, Heeger DJ (2001) Representing retinal image speed in visual cortex. Nat Neurosci 4:461–462 Available at: https://www.nature.com/articles/nn0501_461 [Accessed August 30, 2024].

Sincich LC, Park KF, Wohlgemuth MJ, Horton JC (2004) Bypassing V1: a direct geniculate input to area MT. Nat Neurosci 7:1123–1128 Available at: https://www.nature.com/articles/nn1318 [Accessed September 18, 2024].

Skottun BC, De Valois RL, Grosof DH, Movshon JA, Albrecht DG, Bonds AB (1991) Classifying simple and complex cells on the basis of response modulation. Vision Research 31:1078– 1086 Available at: https://www.sciencedirect.com/science/article/pii/0042698991900332 [Accessed August 30, 2024].

Smith GB, Sederberg A, Elyada YM, Van Hooser SD, Kaschube M, Fitzpatrick D (2015) The development of cortical circuits for motion discrimination. Nat Neurosci 18:252–261.

Snodderly DM, Gur M (1995) Organization of striate cortex of alert, trained monkeys (Macaca fascicularis): ongoing activity, stimulus selectivity, and widths of receptive field activating regions. Journal of Neurophysiology 74:2100–2125 Available at: https://journals.physiology.org/doi/abs/10.1152/jn.1995.74.5.2100 [Accessed June 2, 2021].

Solomon SG, Rosa MGP (2014) A simpler primate brain: the visual system of the marmoset monkey. Front Neural Circuits 8 Available at: https://www.frontiersin.org/journals/neural-circuits/articles/10.3389/fncir.2014.00096/full [Accessed September 18, 2024].

Stacy AK, Schneider NA, Gilman NK, Hooser SDV (2023) Impact of Acute Visual Experience on Development of LGN Receptive Fields in the Ferret. J Neurosci 43:3495–3508 Available at: https://www.jneurosci.org/content/43/19/3495 [Accessed September 3, 2024].

Stitt I, Galindo-Leon E, Pieper F, Engler G, Engel AK (2013) Laminar profile of visual response properties in ferret superior colliculus. Journal of Neurophysiology 110:1333–1345 Available at: https://journals.physiology.org/doi/full/10.1152/jn.00957.2012 [Accessed April 5, 2023].

Stoelzel CR, Bereshpolova Y, Gusev AG, Swadlow HA (2008) The Impact of an LGNd Impulse on the Awake Visual Cortex: Synaptic Dynamics and the Sustained/Transient Distinction. J Neurosci 28:5018–5028 Available at: https://www.jneurosci.org/content/28/19/5018 [Accessed August 30, 2024].

Van Essen DC, Maunsell JHR, Bixby JL (1981) The middle temporal visual area in the macaque: Myeloarchitecture, connections, functional properties and topographic organization. Journal of Comparative Neurology 199:293–326 Available at: https://onlinelibrary.wiley.com/doi/abs/10.1002/cne.901990302 [Accessed August 30, 2024].

Van Hooser SD, Heimel JAF, Nelson SB (2003) Receptive field properties and laminar organization of lateral geniculate nucleus in the gray squirrel (Sciurus carolinensis). J Neurophysiol 90:3398–3418.

van Kerkoerle T, Self MW, Roelfsema PR (2017) Layer-specificity in the effects of attention and working memory on activity in primary visual cortex. Nat Commun 8:13804 Available at: https://www.nature.com/articles/ncomms13804 [Accessed September 18, 2024].

van Kleef JP, Cloherty SL, Ibbotson MR (2010) Complex cell receptive fields: evidence for a hierarchical mechanism. J Physiol 588:3457–3470 Available at: https://www.ncbi.nlm.nih.gov/pmc/articles/PMC2988511/ [Accessed August 30, 2024].

Wang H, Dey O, Lagos W, Callaway EM (2021) Diversity in spatial frequency, temporal frequency, and speed tuning across mouse visual cortical areas and layers. Journal of Comparative Neurology:1–22 Available at: https://onlinelibrary.wiley.com/doi/epdf/10.1002/cne.25404.

Weliky M, Bosking WH, Fitzpatrick D (1996) A systematic map of direction preference in primary visual cortex. Nature 379:725–728 Available at: https://www.nature.com/articles/379725a0 [Accessed June 2, 2021].

Weliky M, Katz LC (1994) Functional mapping of horizontal connections in developing ferret visual cortex: experiments and modeling. J Neurosci 14:7291–7305.

Wypych M, Wang C, Nagy A, Benedek G, Dreher B, Waleszczyk WJ (2012) Standardized F1 – A consistent measure of strength of modulation of visual responses to sine-wave drifting gratings. Vision Research 72:14–33 Available at: https://www.sciencedirect.com/science/article/pii/S0042698912002957 [Accessed August 30, 2024].

Yacoub E, Harel N, Ugurbil K (2008) High-field fMRI unveils orientation columns in humans. Proc Natl Acad Sci U S A 105:10607–10612.

Yacoub E, Shmuel A, Logothetis N, Uğurbil K (2007) Robust detection of ocular dominance columns in humans using Hahn Spin Echo BOLD functional MRI at 7 Tesla. Neuroimage 37:1161–1177.

Yu, H., Farley, B. J., Jin, D. Z., & Sur, M. (2005). The coordinated mapping of visual space and response features in visual cortex. Neuron, 47(2), 267–280. 10.1016/j.neuron.2005.06.011.

Zeki S (2015) Area V5—a microcosm of the visual brain. Front Integr Neurosci 9 Available at: https://www.frontiersin.org/journals/integrative-neuroscience/articles/10.3389/fnint.2015.00021/full [Accessed August 30, 2024].

Zheng A, Scott KE, Schormans AL, Mann R, Allman BL, Schmid S (2023) Differences in Startle and Prepulse Inhibition in Contactin-associated Protein-like 2 Knock-out Rats are Associated with Sex-specific Alterations in Brainstem Neural Activity. Neuroscience 513:96–110 Available at: https://www.sciencedirect.com/science/article/pii/S0306452223000313 [Accessed December 13, 2023].

